# Post-Transcriptional Regulation of Antiviral Gene Expression by *N6*-Methyladenosine

**DOI:** 10.1101/2020.08.05.238337

**Authors:** Michael J. McFadden, Alexa B.R. McIntyre, Haralambos Mourelatos, Nathan S. Abell, Nandan S. Gokhale, Hélène Ipas, Blerta Xhemalçe, Christopher E. Mason, Stacy M. Horner

## Abstract

Type I interferons (IFN) induce hundreds of IFN-stimulated genes (ISGs) in response to viral infection. These ISGs require strict regulation for an efficient and controlled antiviral response, but post-transcriptional controls of these genes have not been well defined. Here, we identify a new role for the RNA base modification *N6*-methyladenosine (m^6^A) in the regulation of ISGs. Using ribosome profiling and quantitative mass spectrometry, coupled with m^6^A-immunoprecipitation and sequencing, we identified a subset of ISGs, including *IFITM1*, whose translation is enhanced by m^6^A and the m^6^A methyltransferase proteins METTL3 and METTL14. We further determined that the m^6^A reader YTHDF1 increases the expression of IFITM1 in an m^6^A binding-dependent manner. Importantly, we found that the m^6^A methyltransferase complex promotes the antiviral activity of type I IFN. Thus, these studies identify m^6^A as a post-transcriptional control of ISG translation during the type I IFN response for antiviral restriction.

## Introduction

The IFN family cytokines are potent inhibitors of viral infection that induce hundreds of ISGs, many of which have antiviral activity (Gonzalez-Navajas et al., 2012; Schoggins and Rice, 2011). Type I IFNs (IFN-α and IFN-β) are produced in response to viral infection, and they activate autocrine and paracrine signaling responses through the JAK-STAT pathway (Stark and Darnell, 2012). Specifically, type I IFNs bind to a dimeric receptor (IFNAR), composed of two subunits, IFNAR1 and IFNAR2. IFNAR engagement then activates the Janus kinases JAK1 and TYK2, which phosphorylate the transcription factors STAT1 and STAT2, inducing their hetero-dimerization and interaction with IRF9, to form the ISGF3 transcription factor complex. ISGF3 then translocates into the nucleus, where it binds to IFN-stimulated response elements within the promoters of ISGs to elicit their transcriptional activation (Stark and Darnell, 2012). Many of these ISGs encode antiviral effector proteins that inhibit multiple stages of viral replication and thus establish an early defense against viral replication (Schoggins, 2019). Dysregulation of type I IFNs can lead to viral susceptibility or autoimmune disease (Banchereau and Pascual, 2006; Teijaro, 2016), demonstrating the importance of tight regulatory control of both IFN activation and the IFN response. Indeed, both activation and suppression of the type I IFN response are coordinated at multiple levels, such as by epigenetic modifiers (Fang et al., 2012; Huang et al., 2002; Liu et al., 2002) or by post-transcriptional mechanisms including microRNA regulation and alternative splicing (Forster et al., 2015; West et al., 2019). However, our overall understanding of post-transcriptional regulation of ISG expression is still emerging. Additionally, while a number of studies have identified subsets of ISGs that have unique transcriptional regulators, other mechanisms that govern the regulation of subclasses of ISGs have not been well characterized (Froggatt et al., 2019; Perwitasari et al., 2011; Seifert et al., 2019).

The RNA base modification m^6^A is deposited on RNA by a methyltransferase complex of METTL3 and METTL14 (METTL3/14), among other proteins (Liu et al., 2014). m^6^A coordinates biological processes through various effects on modified mRNAs (Meyer and Jaffrey, 2017; Shi et al., 2019), including increased mRNA turnover and translation, as well as other processes. These effects are mediated by m^6^A reader proteins, such as YTHDF proteins (Liu et al., 2019a; Wang et al., 2014; Wang et al., 2015). Specifically, YTHDF1 increases translation (Wang et al., 2015), YTHDF2 mediates mRNA degradation (Wang et al., 2014), and YTHDF3 cooperatively enhances both of these processes (Shi et al., 2017), although in some cases these proteins may have overlapping functions (Zaccara and Jaffrey, 2020). Through the actions of m^6^A reader proteins, m^6^A can regulate infection by many viruses through modulation of both viral and host transcripts (Williams et al., 2019). We recently profiled changes to the m^6^A landscape of host mRNAs during *Flaviviridae* infection and identified both proviral and antiviral transcripts regulated by m^6^A during infection (Gokhale et al., 2019). Others have found that m^6^A prevents RNA sensing or regulates the expression of signaling molecules involved in the production of cytokines such as type I IFNs (Chen et al., 2019; Durbin et al., 2016; Feng et al., 2018; Kariko et al., 2005; Lu et al., 2020; Zheng et al., 2017) and that m^6^A can destabilize the *IFNB1* transcript, thereby directly regulating the production of IFN-β (Rubio et al., 2018; Winkler et al., 2019). Therefore, m^6^A plays important roles in viral infection and the antiviral response (Zhang et al., 2019a); however, a role for m^6^A in the response to type I IFN and the production of ISGs has not been described.

Here, we mapped m^6^A in the IFN-β-induced transcriptome and identified many ISGs that are m^6^A-modified. We found that METTL3/14 and m^6^A promote the translation of certain m^6^A-modified ISGs, in part through interactions between the transcripts of m^6^A-modified ISGs and the m^6^A reader protein YTHDF1. Importantly, we found that METTL3/14 and m^6^A-mediated enhancement of ISG expression promotes the antiviral effects of the IFN response, as METTL3/14 perturbation affected the replication of vesicular stomatitis virus (VSV) in IFN-β-primed cells. Together, these results establish m^6^A as a post-transcriptional regulator of ISGs for an effective cellular antiviral response.

## Results

### METTL3/14 regulates the translation of certain ISGs

IFN-β induces the transcription of ISGs to shape the innate response to viral infection (Schoggins and Rice, 2011). To investigate whether m^6^A regulates the type I IFN response, we measured the IFN-β-induced expression of several ISGs with known antiviral functions (Schoggins, 2014) following depletion of the m^6^A methyltransferase complex METTL3/14 in Huh7 cells. The IFN-β-induced protein expression of the ISGs IFITM1 and MX1, but not ISG15 and EIF2AK2 (also called PKR), was reduced following depletion of METTL3/14 (Figure 1A; see Methods for information on IFITM1 antibody specificity). Similar results were also seen following METTL3/14 depletion in A549 cells, primary neonatal human dermal fibroblasts (NHDF), and also at multiple time points (8 h, 16 h, and 24 h) after IFN-β in Huh7 cells (Figure S1A-1C); however we note MX1 protein levels were not as strongly affected in A549 and NHDF cells as in Huh7 cells. Conversely, overexpression of METTL3/14 increased the abundance of IFITM1 and MX1, but not ISG15 and EIF2AK2, in response to IFN-β in Huh7 cells (Figure 1B). Importantly, the METTL3/14-regulated ISGs, IFITM1 and MX1, were not expressed without stimulation of cells by IFN-β (Figure 1A-1B). Therefore, any confounding effects of METTL3/14 perturbation on endogenous IFN-β production are negligible for these experiments.

**Figure 1:**
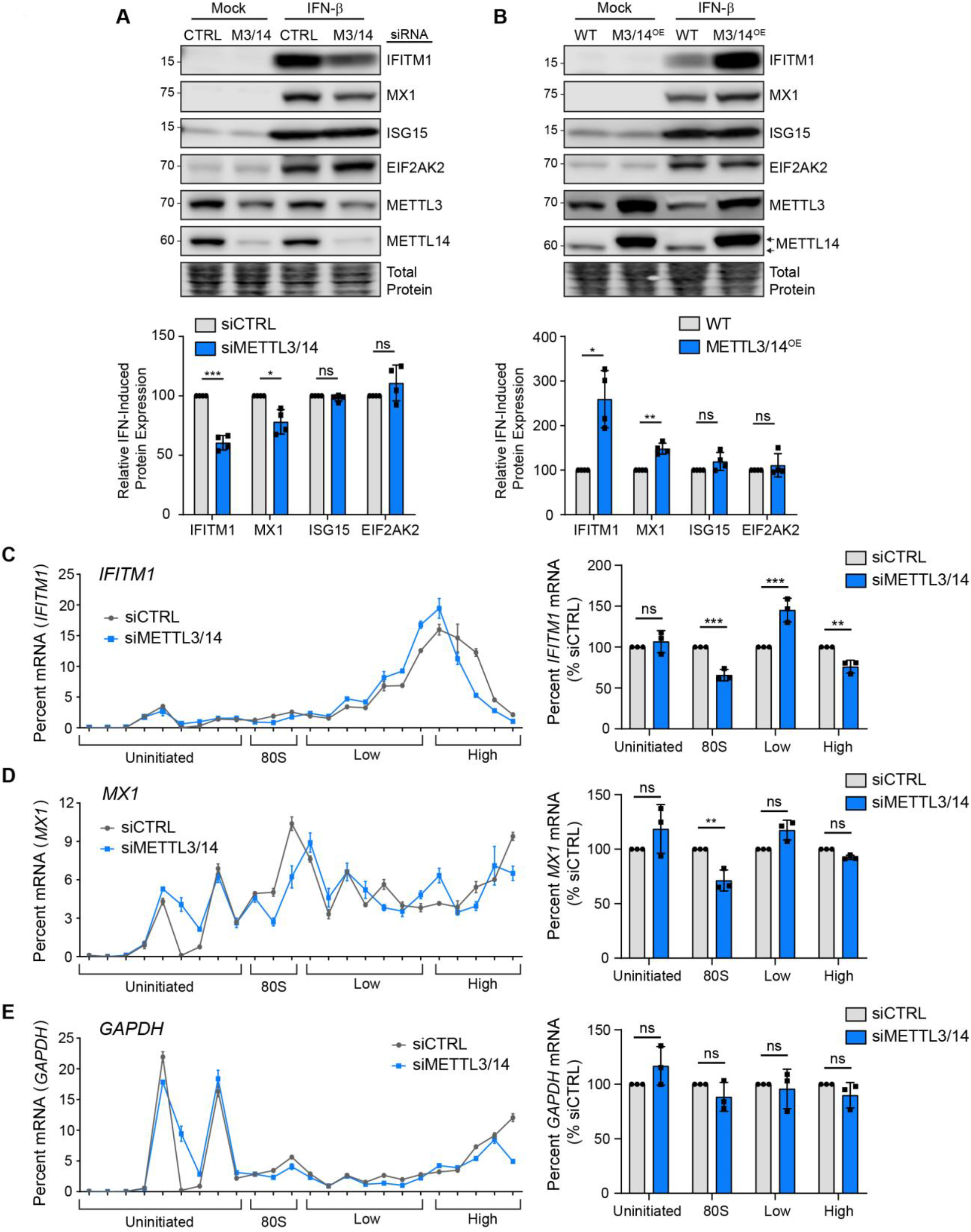
METTL3/14 regulates translation of certain ISGs. **(A, B)** Immunoblot analysis of extracts from Huh7 cells transfected with siRNAs to METTL3/14 (M3/14) or control (CTRL) **(A)** or stably overexpressing FLAG-METTL14 (M3/14^OE^; top arrow denotes FLAG-METTL14; bottom arrow denotes endogenous METTL14) **(B)** prior to mock or IFN-β (24 h) treatment. Relative ISG expression from 4 replicates of (A) and (B) is quantified below relative to siCTRL +IFN-β (A) or WT +IFN-β (B). **(C-E)** RT-qPCR analysis of the relative percentage of *IFITM1* (C), *MX1* (D), and *GAPDH* (E) mRNA across 24 sucrose gradient fractions isolated from extracts of IFN-β-treated (6 h) Huh7 cells treated with CTRL or METTL3/14 siRNA. The uninitiated (free, 40S, and 60S subunits), initiated (80S), low or high molecular weight polysomes, are noted. Graphs on the right show the percentages of mRNAs in combined fractions for *IFITM1*, *MX1*, or *GAPDH*. Percentages from fractions were added to yield the total percentage in each category. Values are the mean ± SEM of 4 biological replicates (A-B), the mean ± SD of 3 technical replicates, representative of 3 experiments (C-E, left graphs), and the mean ± SEM of 3 biological replicates (C-E, right graphs). * p < 0.05, ** p < 0.01, *** p < 0.005 by unpaired Student’s t test (A-B), and 2-way ANOVA with Sidak’s multiple comparisons test (C-E). ns = not significant. See also Figure S1-S2.

The METTL3/14 m^6^A methyltransferase complex regulates many aspects of mRNA metabolism (Liu et al., 2019a). To determine how METTL3/14 regulates the protein abundance of certain ISGs, we first tested whether METTL3/14 depletion led to a decrease in induction of ISG mRNA in response to IFN-β. We measured the induction of ISG mRNA in response to IFN-β over a timecourse using RT-qPCR (Figure S1D). Neither the mRNA abundance nor the kinetics of IFN-β-mediated induction of the METTL3/14-regulated ISGs *IFITM1* and *MX1* were affected by METTL3/14 depletion. The mRNA levels of the non-METTL3/14-regulated ISG *EIF2AK2* was unaffected, while *ISG15* mRNA was increased (Figure S1D). These data indicate that the mRNA abundance of *IFITM1* and *MX1* does not underlie the observed differences in protein levels, suggesting that neither the transcription nor the mRNA stability of these ISGs are regulated by METTL3/14 (Figure S1D). Further, using RNA-seq following IFN-β treatment, we noted little effect of METTL3/14 depletion on the mRNA abundance of a defined set of core ISGs (Shaw et al., 2017) (Figure S1E) or expressed ISGs more broadly (Table S1). These data are in agreement with a previous report that found that collective ISG RNA stability is unaffected by METTL3 depletion (Winkler et al., 2019).

As both METTL3/14 and m^6^A have been shown to promote the nuclear export of certain mRNAs (Lesbirel and Wilson, 2019), we also tested whether the nuclear export of select ISGs was altered by METTL3/14 depletion. However, after IFN stimulation, METTL3/14 depletion did not alter the nuclear-cytoplasmic ratio of the METTL3/14-regulated ISGs *IFITM1* and *MX1*, the non-regulated ISGs *ISG15* and *EIF2AK2*, a non-methylated control *HPRT1* (Wang et al., 2014), or the nuclear-localized control *MALAT1* (Figure S2C). Therefore, METTL3/14 does not regulate these ISGs through changes to their protein stability or nuclear export.

To test whether METTL3/14 regulates the translation of *IFITM1*, we measured the polysome occupancy of *IFITM1* induced by IFN-β in control cells or in those depleted of METTL3/14. METTL3/14 depletion did not change overall polysome density, as observed by the similar relative absorption across fractions (Figure S2D). However, METTL3/14 depletion did result in lower levels of *IFITM1* mRNA in the 80S fractions and a shift from the heavy to the light polysome fractions (Figure 1C), indicating impaired translation of *IFITM1* following METTL3/14 depletion. A similar, yet less pronounced shift was observed for *MX1* (Figure 1D), while the polysome occupancy of the housekeeping control gene *GAPDH* was unaffected (Figure 1E). These results indicate that METTL3/14 regulates the translation of certain ISGs, such as IFITM1 and MX1.

### METTL3/14-regulated ISGs are modified by m^6^A

To determine whether the METTL3/14-regulated ISGs *IFITM1* and *MX1*, as well as other ISGs, are m^6^A-modified, we mapped m^6^A in the IFN-induced transcriptome in Huh7 cells using methylated RNA immunoprecipitation and sequencing (MeRIP-seq) (Dominissini et al., 2012; Meyer et al., 2012). After defining the ISGs in this experiment (Figure S3A; Table S2), we then called peaks in read coverage post-m^6^A immunoprecipitation compared to input using the MeTDiff m^6^A peak caller (Cui et al., 2018) (Table S2). We observed that peaks across mRNAs were enriched around the end of the coding sequence and the beginning of the 3’ UTR, as expected (Dominissini et al., 2012; Meyer et al., 2012) (Figure 2A). The most highly enriched RNA sequence motif within peaks was [U/A]GGAC, which matches the known m^6^A motif of DRA^m^C (Dominissini et al., 2012; Meyer et al., 2012) (Figure 2A). We found that approximately 85% percent of ISGs, classified as those upregulated more than 4-fold following IFN treatment, were m^6^A-modified, as compared to 74% percent of the expressed transcriptome of Huh7 cells (mean coverage ≥ 10) (Figure 2B). This was consistent with a previous study that found that ISGs were m^6^A-modified at a similar percentage to the transcriptome (Winkler et al., 2019). The percent of ISGs that are m^6^A-modified was similar among other classes of ISGs, including a ‘core’ class of ISGs that are evolutionarily conserved among vertebrate species and a subset of 14 of these core ISGs with known antiviral functions (Shaw et al., 2017) (Figure 2B). Plotting the MeRIP-seq reads relative to the input reads of individual genes can be informative of m^6^A status, as m^6^A peak calling methods have known limitations (McIntyre et al., 2020). Thus, we generated plots for *IFITM1*, *MX1*, *ISG15*, and *EIF2AK2* and used the m^6^A peak callers MeTDiff (Cui et al., 2018) and meRIPPer (https://sourceforge.net/projects/meripper/) (Table S2) to reveal that the METTL3/14-regulated genes *IFITM1* and *MX1* had m^6^A peaks (Figure 2C-D), while *ISG15* and *EIF2AK2* lacked called m^6^A peaks (Figure 2E-F). These plots suggested that the 3’ UTR of *ISG15* may also contain an m^6^A site (Figure 2E). We then compared the m^6^A status of ISGs from our MeRIP-seq experiment to data from previously published studies that profiled m^6^A after IFN-inducing treatments, such as dsDNA (Rubio et al., 2018) or human cytomegalovirus (HCMV) infection (Winkler et al., 2019) (Figure S3B). This comparison showed consistent prediction of m^6^A methylation status for core antiviral ISGs among all three studies (Figure S3B). Indeed, dsDNA treatment potently activates IFN production and elicited m^6^A modification of the same core antiviral ISGs found in our experiment. Infection with HCMV also elicited m^6^A modification of certain ISGs, although fewer peaks were called in these ISGs after HCMV infection than after IFN-β treatment or dsDNA treatment (Figure S3B). We note this virus encodes factors to dampen IFN signaling (Miller et al., 1999), therefore ISGs are likely not as strongly induced as compared to dsDNA or IFN-β treatment. The presence of m^6^A on many ISGs suggests that m^6^A may regulate the antiviral type I IFN response.

**Figure 2:**
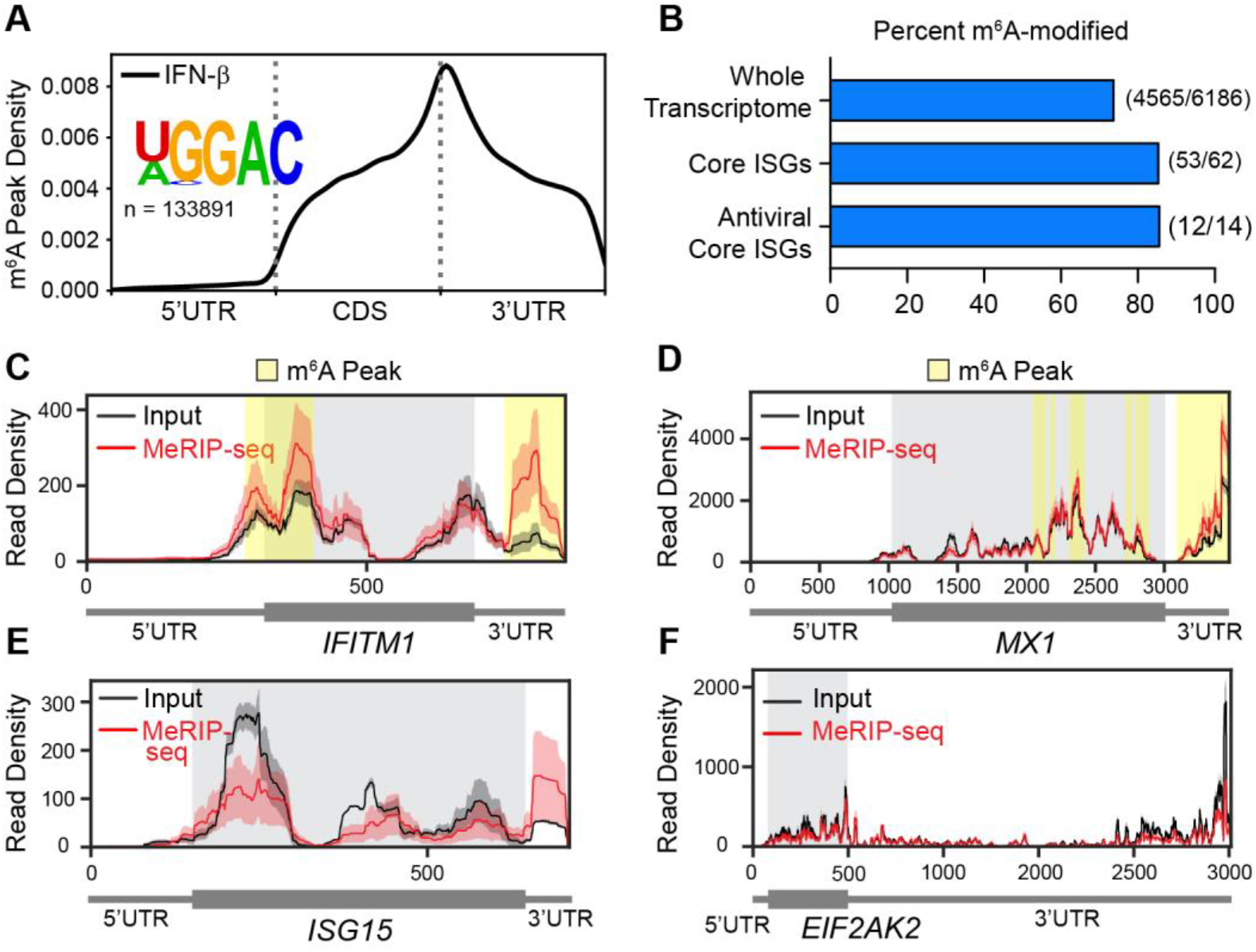
METTL3/14-regulated ISGs are modified by m^6^A. **(A)** Metagene plot of predicted m^6^A distribution across the transcriptome following IFN-β treatment (8 h), with relative positions of DRACH motif sites under statistically significant peaks plotted, as well as the most highly enriched motif under peaks. **(B)** The percent of genes modified by m^6^A in the expressed transcriptome, genes with mRNA induction ≥ 4-fold in response to IFN-β treatment (ISGs), a group of core ISGs conserved in vertebrate species (Shaw et al., 2017), or a subset of these core ISGs with antiviral functions (Shaw et al., 2017). **(C-F)** Read coverage plots of MeRIP (red) and input (black) reads in *IFITM1* (C), *MX1* (D), *ISG15* (E), and *EIF2AK2* (F) transcripts. Variance between biological replicates is represented by red and black shading around read coverage. Gray shading represents coding sequence, yellow shading represents m^6^A peaks called by MeTDiff (Cui et al., 2018) and meRIPPer (https://sourceforge.net/projects/meripper/) software. All analyses are performed on 3 biological replicates. See also Figure S3.

### m^6^A modification in the 3’ UTR of IFITM1 enhances its translation

m^6^A is known to enhance the translation of certain mRNAs (Coots et al., 2017; Gokhale et al., 2019; Lin et al., 2016; Mao et al., 2019; Wang et al., 2015). Specifically, the m^6^A reader protein YTHDF1 can recognize m^6^A within 3’ UTRs and associate with eukaryotic translation initiation factors such as eIF3 to enhance the translation of m^6^A-modified transcripts (Wang et al., 2015). To determine whether the translational regulation of ISGs by METTL3/14 is elicited through m^6^A, we used *IFITM1* as a model METTL3/14-regulated ISG. We first determined the effect of METTL3/14 depletion on m^6^A modification of *IFITM1*. MeRIP-RT-qPCR revealed that *IFITM1* mRNA was enriched above the m^6^A-negative ISG *EIF2AK2* and a spiked-in m^6^A-negative synthetic RNA, confirming that it contains m^6^A. METTL3/14 depletion reduced the m^6^A-enrichment of *IFITM1* mRNA but not of the m^6^A-negative *EIF2AK2* transcript or the m^6^A-negative synthetic RNA (Figure 3A-B). These data reveal that *IFITM1* is m^6^A-modified by METTL3/14.

**Figure 3:**
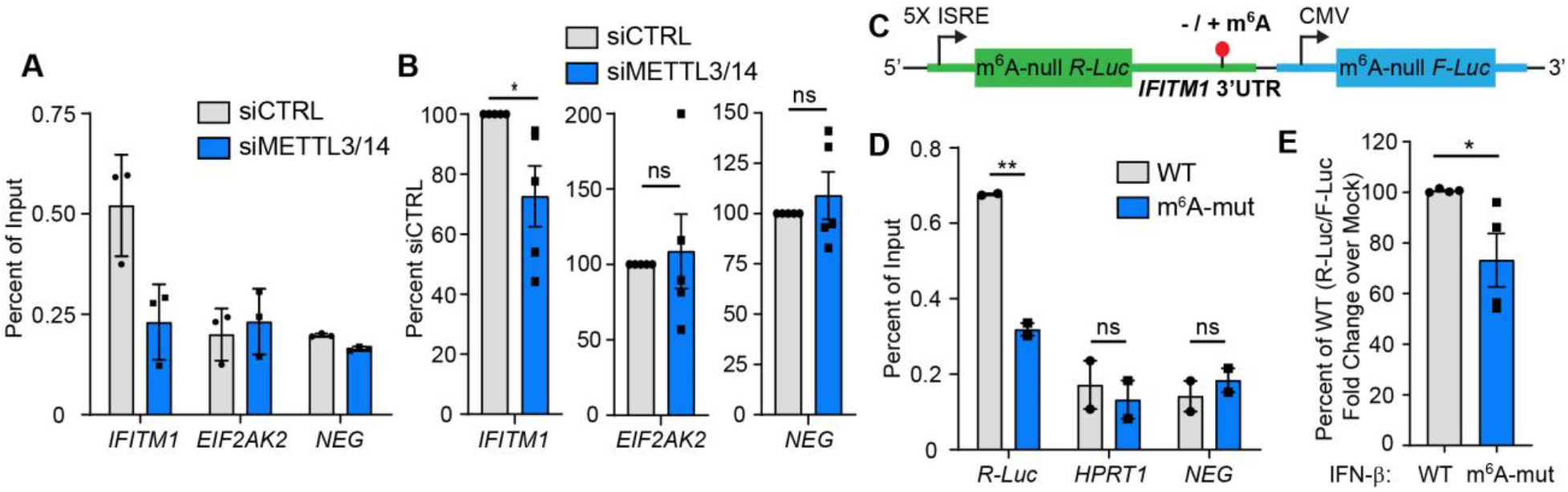
m^6^A modification of the *IFITM1* 3’ UTR enhances translation. **(A)** Representative MeRIP-RT-qPCR analysis of relative m^6^A level of *ISGs* induced by IFN-β (8 h) in Huh7 cells treated with non-targeting control (siCTRL) or METTL3/14 siRNA and spiked-in m^6^A-negative (NEG) oligonucleotides. **(B)** Relative percent enrichment of each gene in (A), normalized to siCTRL, from 5 biological replicates. **(C)** Schematic of WT and mutant ISRE-m^6^A-null *Renilla* luciferase (R-Luc) *IFITM1* 3’ UTR reporters that also express m^6^A-null firefly luciferase (F-Luc) from a separate promoter. **(D)** MeRIP-RT-qPCR analysis of relative m^6^A level of WT and m^6^A-mut *IFITM1* 3’ UTR reporter RNA from transfected Huh7 cells treated with IFN-β (8 h). **(E)** Relative luciferase activity (R-Luc/F-Luc) in IFN-β induced (8 h, relative to mock) WT and m^6^A-mut *IFITM1* 3’ UTR reporters. Values are the mean ± SD of 3 technical replicates representative of 5 biological replicates (A); the mean ± SEM of 5 biological replicates (B); the mean ± SEM of 2 biological replicates (D), or mean ± SEM of 4 biological replicates (E). * p < 0.05, ** p < 0.01 by unpaired Student’s t test. ns = not significant.

Having confirmed that *IFITM1* is m^6^A modified, we next generated a luciferase reporter that contains an IFN-stimulated response element (ISRE) promoter-driven *Renilla* luciferase in which all DRAC motifs were ablated (m^6^A-null *R-Luc*) (Gokhale et al., 2019) fused to the wild type (WT) *IFITM1* 3’ UTR, or an analogous 3’ UTR sequence in which the four putative m^6^A motifs under the m^6^A peak in the 3’ UTR in *IFITM1* were inactivated by A→G transitions (m^6^A-mut) (Figure 3C). These constructs also express a CMV promoter-driven m^6^A-null firefly luciferase gene as a control. The m^6^A modification status of the *IFITM1* 3’ UTR reporter was first assessed using MeRIP-RT-qPCR after IFN-β treatment. The WT *IFITM1* 3’ UTR reporter had increased m^6^A modification compared to the m^6^A-mut *IFITM1* 3’ UTR reporter, as well as the negative controls: *HPRT1*, which does not contain m^6^A (Wang et al., 2014), and an m^6^A-negative synthetic RNA control (Figure 3D). We next compared the production of the *Renilla* luciferase protein, relative to firefly luciferase, from the WT and m^6^A-mut IFITM1 3’ UTR reporters by measuring luciferase activity. We found that the relative luciferase activity of the m^6^A-mut IFITM1 3’UTR reporter was significantly decreased following IFN-β treatment compared to the WT IFITM1 3’ UTR reporter (Figure 3E). Together, these data suggest that METTL3/14 regulates *IFITM1* translation through addition of m^6^A to the 3’ UTR and that m^6^A within the *IFITM1* 3’ UTR is sufficient to enhance its translation.

### YTHDF1 enhances IFITM1 protein expression in an m^6^A-dependent fashion

The m^6^A binding protein YTHDF1 enhances translation of a number of m^6^A-modified genes (Wang et al., 2015). To test if YTHDF1 elicited the translation-promoting effects of m^6^A on ISGs, we stably overexpressed YTHDF1 (Y1^OE^) or an m^6^A binding-deficient YTHDF1 protein (Xu et al., 2015) (Y1mut^OE^) in Huh7 cells and measured the IFN-induced expression of ISGs 24 hours later. Overexpression of YTHDF1 was sufficient to increase IFITM1 protein expression in response to IFN-β, while overexpression of the m^6^A binding-deficient YTHDF1 protein (Y1mut^OE^) did not increase IFITM1 abundance (Figure 4A-B). Importantly, wild-type and mutant YTHDF1 overexpression did not significantly affect the levels of *IFITM1* mRNA following IFN-β treatment, suggesting that YTHDF1 does not directly regulate IFN signaling or *IFITM1* mRNA stability (Figure 4C). Neither the IFN-induced expression of the m^6^A-containing ISG MX1, nor the non-m^6^A containing ISGs ISG15 and EIF2AK2, were significantly altered by YTHDF1 overexpression (Figure 4A-B). Interestingly, we found that WT YTHDF1 bound to the transcripts of *IFITM1*, *MX1*, *ISG15*, and the m^6^A*-*positive control *SON* (Wang et al., 2014), while the m^6^A-binding-defective YTHDF1 mutant protein did not. The non-m^6^A containing mRNAs *EIF2AK2* and *RPL30* (Wang et al., 2015) did not bind to either protein (Figure 4D-E). Together, these results reveal that YTHDF1 binds to m^6^A on *IFITM1* and enhances its translation. The apparent m^6^A-dependent binding of YTHDF1 to *ISG15* suggests that *ISG15* is actually m^6^A-modified. In fact, plotting MeRIP-seq reads over input reads for *ISG15* did show a potential region of m^6^A enrichment in its 3’ UTR (Figure 2E), although this was not identified as significant by two peak callers (Table S2).Taken together, these data suggest that YTHDF1 has transcript-specific roles in promoting translation, as it bound *IFITM1*, *MX1*, and *ISG15*, but its overexpression was only sufficient to significantly increase the protein production of IFITM1.

**Figure 4:**
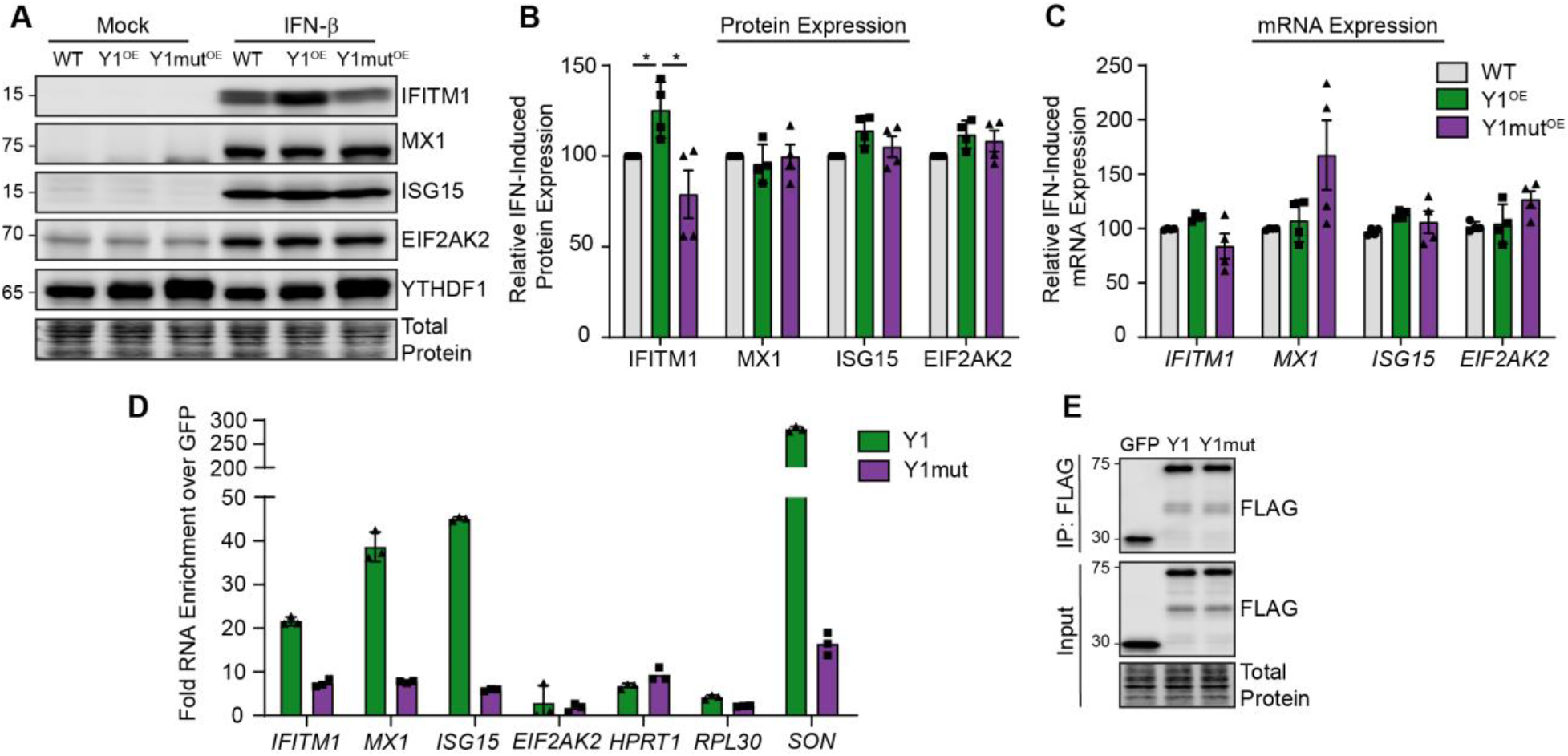
YTHDF1 enhances IFITM1 protein expression in an m^6^A-dependent fashion. **(A)** Immunoblot analysis of extracts from Huh7 cells stably overexpressing FLAG-YTHDF1 WT (Y1^OE^) or FLAG-YTHDF1 W465A (Xu et al., 2015) (Y1mut^OE^) following mock or IFN-β (24 h) treatment. **(B)** Quantification of ISG expression following IFN-β from 3 independent experiments of (A), normalized to total protein and graphed relative to siCTRL. **(C)** RT-qPCR analysis of ISG mRNA expression normalized to HPRT1 in Huh7 cells stably overexpressing FLAG-YTHDF1 WT (Y1^OE^) or W465A (Y1mut^OE^) after IFN-β (24 h) treatment **(D)** RT-qPCR analysis of enrichment of mRNAs following immunoprecipitation of FLAG-YTHDF1 WT (Y1) or W465A (Mut) compared to FLAG-GFP from Huh7 cells following IFN-β (8 h). IP values are normalized to input values and plotted as fold enrichment over GFP. **(E)** Immunoblot of FLAG-immunoprecipitated and input fractions used in (D). Values in (B-C) are the mean ± SEM of 3 biological replicates. * p < 0.05, by Kruskal-Wallis with Dunn’s multiple comparisons test. Everything unlabeled was not significant with p > 0.05. Values in (D) are the mean ± SD of 3 technical replicates and are representative of 4 independent experiments.

### METTL3/14 and m^6^A promote the translation of a subset of ISGs

Having demonstrated that m^6^A supports the translation of two ISGs (*IFITM1* and *MX1*), and that m^6^A is present on many ISGs, we next sought to identify additional ISGs whose protein expression is regulated by METTL3/14. To this end, we employed quantitative mass spectrometry-based proteomics with stable isotope labeling of amino acids (SILAC) to compare the proteomes of siCTRL and siMETTL3/14 cells after IFN-β treatment (Table S3). The effect of siMETTL3/14 compared to siCTRL on protein abundance is centered at a log ratio of 0 for the majority of proteins (Figure S4A), demonstrating that METTL3/14 depletion does not have a global effect on protein levels after IFN-β treatment. We determined which proteins are ISGs by defining ISGs as genes upregulated >2-fold by IFN-β treatment in our previous RNA-seq experiment (Table S1). While mass spectrometry detection of ISGs was limited (n=18), we did identify a number of METTL3/14-regulated ISGs (Figure 5A, MS). The protein expression of most of these ISGs was decreased following METTL3/14 depletion, and these ISGs included the previously identified m^6^A-modified IFITM1 (peptides corresponding to IFITM1/2/3) and MX1, as well as additional antiviral ISGs such as OAS2 and the different HLA-C chains (Figure 5A), which are also m^6^A-modified. We also compared these data to our previous RNA-seq experiment (Table S1) to determine whether the effects of METTL3/14 on the protein level of these ISGs is determined by regulation of their mRNA expression. Importantly, following METTL3/14 depletion, the ISGs in this experiment that were decreased at the protein level did not also have a decrease in mRNA abundance, suggesting they may be regulated at the translation level, as our earlier polysome profiling indicated for IFITM1 and MX1 (Figure 1C-D; 5A, RNA). We note that not all m^6^A-modified ISGs identified by mass spectrometry were regulated by METTL3/14 depletion (Figure 5A, m^6^A). This suggests that METTL3/14 and m^6^A regulate a subset of ISGs and support their protein expression.

**Figure 5:**
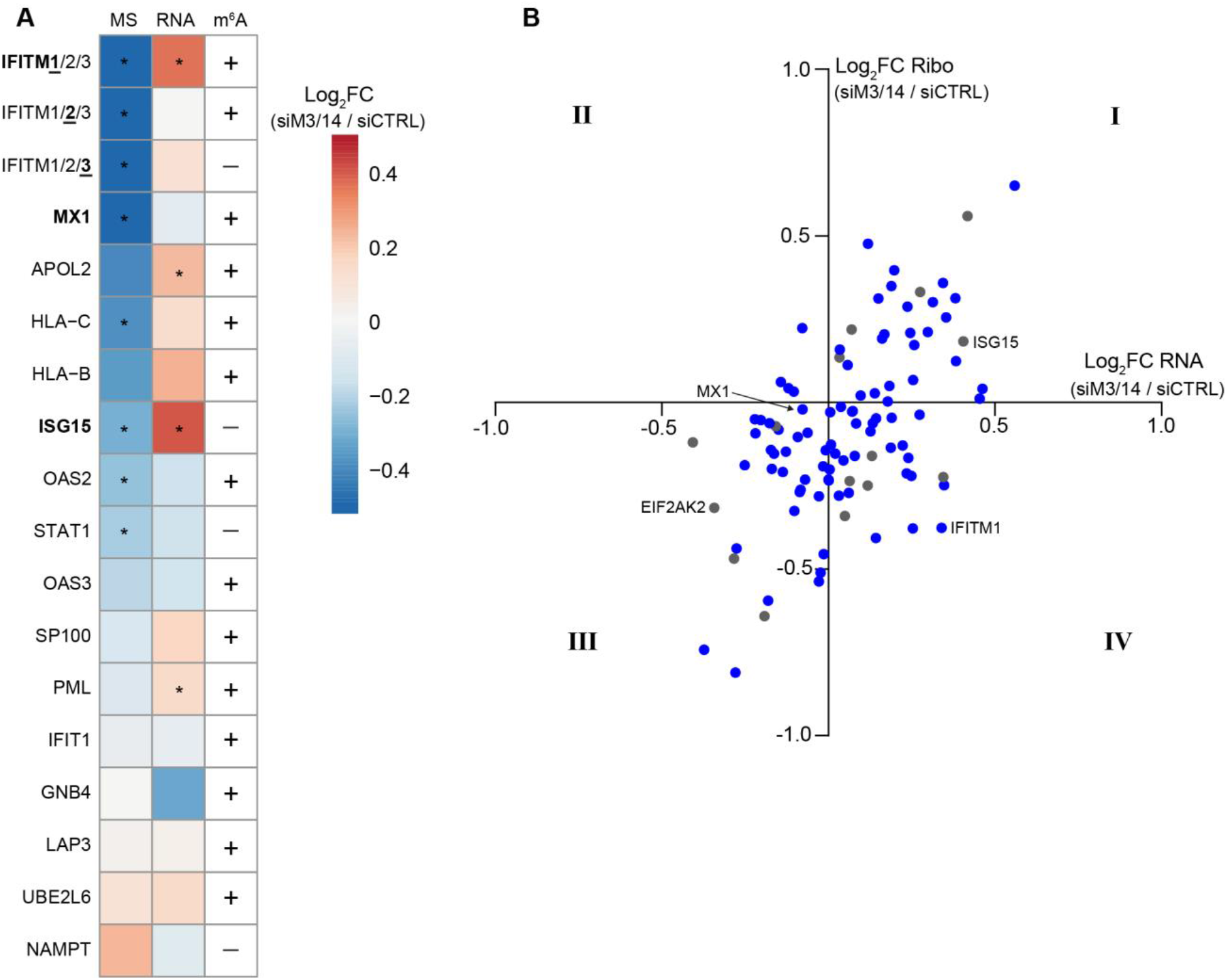
METTL3/14 regulates the translation of a subset of ISGs. **(A)** 3-column heatmap shows the effect of METTL3/14 depletion on the expression of ISGs in Huh7 cells following IFN-β treatment. The first column shows the log_2_ fold change of protein estimates from quantitative mass spectrometry (siMETTL3/14 over siCTRL + IFN-β 24 h; n=2 biological replicates). The second column shows log_2_ fold change of mRNA reads from an independent RNA-seq experiment (siMETTL3/14 over siCTRL + IFN-β 8 h; n=3 biological replicates), and the third column indicates m^6^A status (+ indicates m^6^A-positive; - indicates m^6^A-negative) from MeRIP-seq (+ IFN-β 8 h; n=3 biological replicates). Genes include any ISGs induced more than 2-fold by IFN from RNA-seq that were also detected by mass spectrometry. ISGs investigated in other figures are shown in bold. Because IFITM1/2/3 are similar, we used this notation to indicate peptides detected from this family of proteins; however, RNA-seq fold change and m^6^A status correspond to the underlined number. * adjusted P < 0.05. **(B)** Four-quadrant scatterplot showing the effect of METTL3/14 on the expression of ISGs. The y-axis is the log_2_ fold change of ribosome protected fragments from Ribo-seq (siMETTL3/14 over siCTRL), and the x-axis is the log_2_ fold change of mRNA reads from an independent RNA-seq experiment (siMETTL3/14 over siCTRL). m^6^A-modified (blue) or m^6^A-negative (gray) genes are noted. ISGs investigated in other figures are labeled. See also Figure S4.

As a complementary approach, we also used Ribo-seq to more broadly define the role of METTL3/14 in translational regulation of ISGs (Table S4). As ribosome profiling relies on digestion of mRNA that is not ribosome-bound, we first confirmed that reads in the untranslated regions were depleted (Figure S4B). Then we analyzed the top 100 most highly-induced ISGs (Table S1) that were actively translated (base mean >25) and compared the effect of METTL3/14 depletion on ribosome density (Ribo) to mRNA abundance from our previous RNA-seq analysis (RNA) (Figure 5B; Figure S4C; Table S1). METTL3/14 depletion overall appeared to result in decreased ribosome occupancy among many of these ISGs (66/100, including *IFITM1*), without having any generalized effect on their mRNA abundance (Figure S4C). In many cases, METTL3/14 depletion affected both the mRNA abundance and ribosome protection of individual ISGs similarly (Figure 5B, Quadrants I and III). However, for roughly a third of these genes (33/100), METTL3/14 depletion resulted in decreased ribosome protection, despite greater mRNA abundance (Figure 5B, Quadrant IV). Alternatively, very few (4/100) ISGs had both increased ribosome protection and decreased mRNA abundance following METTL3/14 depletion (Figure 5B, Quadrant II). We note that, of these 100 ISGs, 85 were m^6^A-modified, roughly consistent with the 74% of genes that we had identified in the total expressed transcriptome as containing m^6^A (Table S2). Interestingly, a number of m^6^A-modified ISGs were not regulated by METTL3/14, as measured by ribosome protection or mRNA abundance, supporting a role for METTL3/14 and m^6^A in regulation of only certain ISGs. These data, taken together with our quantitative mass spectrometry and RNA-seq analysis, suggest that METTL3/14 regulates the translation of a subset of ISGs to support their protein expression during the type I IFN response.

### METTL3/14 augments the antiviral effects of the IFN response

The fact that METTL3/14 enhances the expression of a subset of ISGs during the type I IFN response suggests that METTL3/14 could be required for an optimal antiviral response. To determine whether METTL3/14 contributes to viral restriction by type I IFN, we measured the ability of type I IFN to restrict infection by the negative-sense stranded RNA virus, vesicular stomatitis virus (VSV), following METTL3/14 perturbation. The VSV genome contains the m^6^Am cap modification, but as the deposition of this modification is not controlled by METTL3/14 (Boulias et al., 2019; Ogino and Banerjee, 2011; Sendinc et al., 2019), we would not expect VSV replication to be directly affected by changes in the levels of METTL3/14. Rather, any impact on VSV replication would likely be mediated by methylation of host factors. We perturbed the expression of METTL3/14 using siRNAs or by overexpression and then determined the percent of cells infected by VSV at 6 hours post-infection in the presence and absence of a low dose of IFN-β pretreatment (6 hours; 25 U/mL). Measuring VSV infection at early time points after infection allowed us to measure viral replication prior to cellular upregulation of ISGs induced directly in response to infection. Indeed, in the absence of IFN-β pretreatment, we saw no induction of ISGs by VSV in any condition (Figure 6A-B). Additionally, as anticipated, we found that the replication of VSV, as measured by immunoblotting or quantifying the percent of cells infected, was not altered by depletion or overexpression of METTL3/14 in cells in the absence of IFN-β pretreatment (Figure 6). As observed earlier, following IFN-β pretreatment, METTL3/14 depletion led to decreased expression of METTL3/14-regulated ISGs (Figure 6A), while METTL3/14 overexpression increased the expression of these ISGs (Figure 6B). As expected, IFN-β pretreatment resulted in overall less replication of VSV, as IFN-β is known to inhibit VSV replication (Muller et al., 1994) (Figure 6). However, upon depletion of METTL3/14, the ability of IFN-β to restrict VSV was reduced (Figure 6A, 6C). Conversely, METTL3/14 overexpression enhanced IFN-mediated restriction of VSV (Figure 6B, 6D). These data indicate that METTL3/14 enhances the antiviral properties of type I IFN and is required for an efficient IFN-mediated antiviral response.

**Figure 6:**
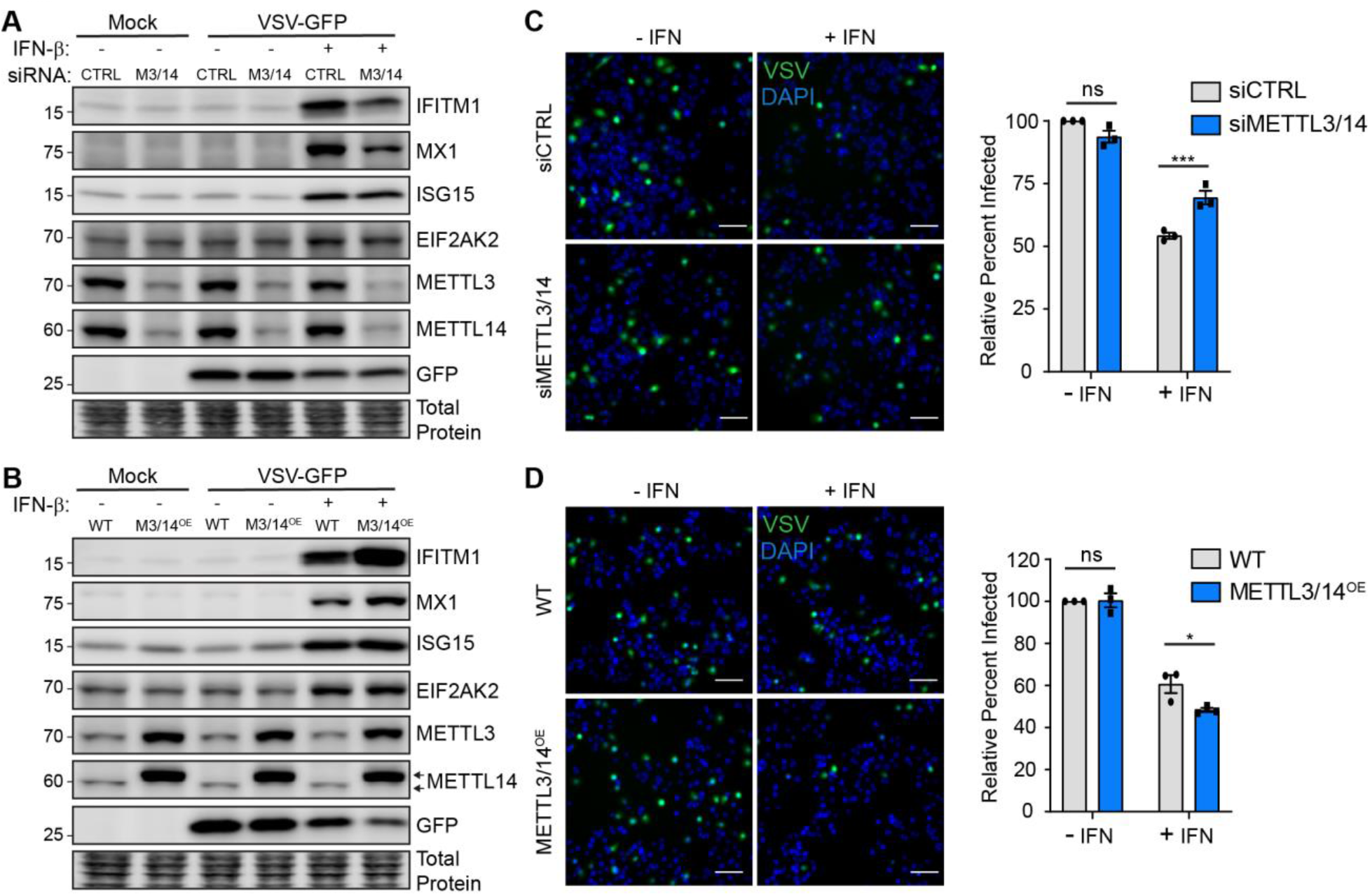
METTL3/14 augments the antiviral effects of the type I IFN response. **(A, B)** Representative immunoblot analysis (n=3) of extracts from Huh7 cells transfected with siRNAs or stably overexpressing FLAG-METTL14, which also enhances METTL3 expression (M3/14^OE^); (B), then treated with IFN-β (6 h) or mock, followed by infection with VSV (MOI=2; 6 h). Arrows denote FLAG-METTL14 (top) and endogenous METLL14 (bottom). **(C, D)** Representative micrographs of Huh7 cells treated with non-targeting control (siCTRL) or METTL3/14 siRNA (C) or stably overexpressing FLAG-METTL14 (METTL3/14^OE^; D), that were pre-treated with IFN-β (6 h), and then infected with VSV (MOI=2; 6 h), with quantification of percent of cells infected from 3 independent experiments with 5 fields per condition, with >150 cells per field, normalized to siCTRL or WT with no IFN treatment, shown on the right. Scale Bar, 100 μm. Values are the mean ± SEM of 3 biological replicates. * p < 0.05, *** p < 0.001 by 2-way ANOVA with Sidak’s multiple comparisons test. ns = not significant.

## Discussion

Post-transcriptional control of the type I IFN response remains poorly understood, and most of our existing knowledge centers around miRNA-mediated regulation of the IFN-induced JAK-STAT signaling pathway (Forster et al., 2015) or a few examples of alternative splicing of ISG transcripts (West et al., 2019). While a number of reports have documented non-canonical activation or delayed stimulation of subsets of ISGs during viral infection, the molecular pathways that can control these subsets of ISGs are not well understood (Pulit-Penaloza et al., 2012; Rose et al., 2010). However, a number of studies have identified transcriptional regulators of subsets of ISGs (Froggatt et al., 2019; Perwitasari et al., 2011; Seifert et al., 2019), and the mRNA Cap1 methyltransferase CMTR1 was recently shown to regulate the expression of certain ISGs (Williams et al., 2020). These studies demonstrate the complexity of regulation of expression of ISGs that extends beyond transcriptional induction of ISGs from signaling of the JAK-STAT pathway. Here, we identify a novel post-transcriptional regulatory mechanism for the expression of a subset of antiviral ISGs. We found that the m^6^A methyltransferase complex of METTL3/14 methylates certain antiviral ISGs to facilitate their translation to promote an antiviral cellular state.

The transcript-specific effects of m^6^A can modulate gene expression to coordinate cellular responses. Indeed, we found that the presence of m^6^A on ISGs can elicit different mechanisms of post-transcriptional regulation. For *IFITM1*, m^6^A in the 3’ UTR led to an increase in its translation, via METTL3/14 and the reader protein YTHDF1. Consistent with our results, previous reports have shown that 3’ UTR m^6^A modification enhances translation initiation and that YTHDF1 likely mediates this enhancement by recruiting eIF3 to m^6^A-modified mRNAs (Wang et al., 2015). Interestingly, while the m^6^A-modified *MX1* is also upregulated at the protein level by METTL3/14, YTHDF1 overexpression was not sufficient to elicit this upregulation. This may indicate that MX1 requires other factors or additional readers to enhance its expression. Indeed, YTHDF3 has recently been shown to have roles in promoting translation of m^6^A-modified genes, perhaps by its interaction with proteins of the 40S and 60S ribosomal subunits (Li et al., 2017; Shi et al., 2017), and YTHDC2 can recognize m^6^A within coding sequence to enhance translation (Mao et al., 2019). Of note, others have found that YTHDF3 inhibits ISG production in murine models through its enhancement of *FOXO3* translation, although this apparently occurred independently of m^6^A (Zhang et al., 2019b). Therefore, m^6^A and its related proteins can regulate ISG expression through a variety of mechanisms. Indeed, only a subset of our identified m^6^A-modified ISGs were translationally enhanced by METTL3/14, as shown by a combination of Ribo-seq, quantitative mass spectrometry, and RNA-seq (Figure 5). As m^6^A has multiple functions in mRNA metabolism, it is possible that m^6^A affects processes other than translation for these other modified ISGs, for example by modulating their splicing, nuclear export, secondary structure, or stability (Liu et al., 2019a). Indeed, it is likely that *ISG15* mRNA stability is regulated by m^6^A, as we found that this transcript is bound by YTHDF1, appears to have an m^6^A site in its 3’ UTR, and its mRNA levels are increased following METTL3/14 depletion. m^6^A may also regulate mRNA trafficking or turnover of ISGs at later timepoints after IFN stimulation or may contribute to alternative splicing of antiviral genes in response to IFN.

Disentangling the regulatory effects of m^6^A on viral infection has been challenging, as both viral and host transcripts contain m^6^A (Williams et al., 2019). Recent work by our group and others revealed that m^6^A regulates several aspects of the host response to infection (Gokhale et al., 2019; Rubio et al., 2018; Winkler et al., 2019). For example, when the *IFNB1* transcript is induced, which can occur in response to viral infection, it is modified by m^6^A, and this destabilizes the transcript. This regulation of *IFNB1* may serve as an intrinsic mechanism to dampen and control the innate immune response (Rubio et al., 2018; Winkler et al., 2019). Interestingly, HCMV appears to hijack this arm of immune regulation by upregulating METTL3/14 expression to increase m^6^A on *IFNB1*, which ultimately decreases IFN-β production, resulting in enhanced viral replication (Rubio et al., 2018; Winkler et al., 2019). However, in our work, by directly stimulating ISGs with IFN-β, we reveal additional m^6^A-mediated regulation of certain ISGs downstream of IFN-β production. Specifically, we show that METTL3/14 depletion reduces the ability of IFN to restrict VSV infection, while METTL3/14 overexpression enhances the ability of IFN to inhibit VSV infection (Figure 6). Importantly, as VSV replication was not affected by changes in METTL3/14 expression in the absence of IFN, this suggests that the differential ability of IFN to restrict VSV following perturbation of METTL3/14 expression is not mediated by direct regulation of the viral RNA (Figure 6). Rather, these data support the idea that METTL3/14 augments the antiviral response by enhancing the production of ISGs. Identifying the factors that control m^6^A addition to a specific subset of ISGs will be an important future pursuit and may clarify why only a subset of these antiviral genes become methylated. Many type I IFN-stimulated genes are also induced by type II (IFN-γ) and type III (IFN-λ) IFNs. Future studies may uncover whether signaling downstream of these IFN families also leads to similar m^6^A-mediated modulation of ISG expression. Additionally, exploring whether viruses employ strategies to counter METTL3/14-mediated enhancement of ISGs will shed further light on the interplay between viral and host RNA processes and how RNA modifications regulate these processes.

In addition to regulating type I IFN pathways, m^6^A tunes other cellular responses to viral infection. We recently showed changes to the m^6^A status of certain host transcripts in response to infection by *Flaviviridae*. Further, we found that many of these m^6^A-altered genes regulate *Flaviviridae* infection (Gokhale et al., 2019). Some of the alterations in m^6^A during infection were driven by innate immune sensing pathways, revealing that innate immune activation can affect cellular m^6^A distribution during infection. Others have recently shown that VSV infection impairs the demethylase activity of ALKBH5, resulting in increased m^6^A modification and destabilization of the *OGDH* transcript. This resulted in less production of the metabolite itaconate, which appeared to be required for VSV replication (Liu et al., 2019b). While these effects of m^6^A on VSV replication occurred independently of IFN signaling, our work revealed that m^6^A can also inhibit VSV replication by promoting ISG expression during IFN signaling. While we did not find an effect of m^6^A on VSV replication in the absence of IFN signaling, as described above (Liu et al., 2019b), we did not investigate a role for ALKBH5. Taken together, our findings add to the knowledge of the diverse regulatory functions of m^6^A during host-pathogen interactions.

In summary, we reveal a subset of ISGs that are post-transcriptionally regulated by METTL3/14 through m^6^A modification. Additionally, we show that the translation of these ISGs is enhanced by m^6^A and postulate that m^6^A may be utilized during the IFN response as a strategy for efficient production of antiviral proteins and the establishment of an antiviral cellular state. Together, these data provide a new molecular understanding of type I IFN response regulation that will ultimately broaden our understanding of innate immunity and host-pathogen interactions. In addition to their functions in antiviral innate immunity, ISGs are also known to regulate inflammation and cell death and recent reports have discovered roles for ISGs in cancer and embryonic development (Buchrieser et al., 2019; Cheon et al., 2014; Yockey and Iwasaki, 2018). Therefore, characterizing the molecular mechanisms that govern ISG expression will be essential for understanding their dysregulation and this information could be harnessed to develop therapeutics to alter ISG expression, which will be relevant to multiple diseases.

## Acknowledgements

We thank colleagues who provided reagents (see Methods), New England Biolabs for the gift of anti-m^6^A antibodies, the Duke Functional Genomics Core Facility, the Duke Center for Genomic and Computational Biology Core, the Epigenomics Core and the Scientific Computing Unit at Weill Cornell, and Horner lab members for manuscript discussion. This work was supported by funds from Burroughs Wellcome Fund (S.M.H.) and National Institutes of Health: R01AI125416, R21AI129851 (S.M.H. and C.E.M.), R01MH117406 (C.E.M.), T32-CA009111 (M.J.M.), R01-GM127802 (B.X.). Other funding sources include: National Science and Engineering Research Council of Canada (A.B.R.M. PGS-D funding), American Heart Association (N.S.G. Pre-doctoral Fellowship, 17PRE33670017), National Institute of General Medical Sciences of the National Institutes of Health Medical Scientist Training Program grant to the Weill Cornell/Rockefeller/Sloan Kettering Tri-Institutional MD-PhD Program. (H.M. T32GM007739), Bert L. and N. Kuggie Vallee Foundation, WorldQuant Foundation, Pershing Square Sohn Cancer Research Alliance, NASA (NNX14AH50G).

## Author contributions

Conceptualization: M.J.M. and S.M.H. Investigation: M.J.M., N.S.G., and H.I. Formal analysis: M.J.M., A.B.R.M, H.M., N.S.A., and. Software: A.B.R.M., H.M., N.S.A. Writing – original draft: M.J.M. and S.M.H. Writing – review and editing: M.J.M., A.B.R.M., H.M., N.S.A., N.S.G., B.X., C.E.M., and S.M.H. Funding acquisition: M.J.M., A.B.R.M., N.S.G., B.X., C.E.M., and S.M.H.

## Methods

### Cell Lines

Human hepatoma Huh7 cells, lung carcinoma A549 cells, neonatal human dermal fibroblast (NHDF) cells, Vero cells, and embryonic kidney 293T cells were grown in Dulbecco’s modification of Eagle’s medium (DMEM; Mediatech) supplemented with 10% fetal bovine serum (Thermo Fisher Scientific), 1X minimum essential medium non-essential amino acids (Thermo Fisher Scientific), and 25 mM HEPES (Thermo Fisher Scientific) (cDMEM). The identity of the Huh7 cells used in this study was verified by using the GenePrint STR kit (Promega) (DNA Analysis Facility, Duke University, Durham, NC, USA). A549 cells, 293T, and Vero cells (CCL-185, CRL-3216, and CCL-81) were obtained from American Type Culture Collection (ATCC), NHDF cells (CC-2509) were obtained from Lonza, and Huh7 cells were a gift of Dr. Michael Gale. All cell lines were verified as mycoplasma free by the LookOut Mycoplasma PCR detection kit (Sigma).

### IFN-β Treatment

All IFN-β (PBL Assay Science) treatments were performed at a concentration of 50 units/mL in cDMEM, unless otherwise noted.

### VSV Infection

GFP-expressing VSV (Whelan et al., 2000) was obtained from Dr. Sean Whelan and propagated by infecting Vero cells grown in cDMEM for 48 hours, after which infectious supernatant was harvested and cleared by centrifugation (1,000 X g for 10 minutes at 4°C) and frozen at −80°C prior to titering. To determine the titer of viral stocks, confluent Vero cells were inoculated with serial dilutions of VSV in serum-free DMEM for 2 hours, overlaid with cDMEM containing 2% SeaPlaque Agarose (Lonza), and incubated at 37°C for an additional 24 hours. Cells were then fixed using 4% formaldehyde and visualized to count GFP-expressing plaques and calculate plaque forming units/mL. Experimental VSV infections were performed at a multiplicity of infection of 2 in serum-free DMEM for 1.5 h, after which cDMEM was replenished. Cells were fixed in 4% formaldehyde, washed with PBS, and stained for DAPI (4′,6-diamidino-2-phenylindole) (Life Technologies, 1:1000). For each condition, 5 images were acquired at 10X magnification on a Zeiss Axio Observer Z1 microscope, and images were processed using ZEN 2 (Zeiss). The percent of cells infected was calculated by counting the number of GFP-positive cells / the number of nuclei (DAPI).

### Plasmids

These plasmids have been described previously: pLEX-FLAG-YTHDF1 (Kennedy et al., 2016), psiCheck2-m^6^A-null (Gokhale et al., 2019), psPAX2 (Addgene plasmid #12260; RRID:Addgene_12260), and pMD2.G (Addgene plasmid # 12259; RRID:Addgene_12259). The following plasmids were constructed in this study: pLEX-FLAG-METTL14, pLEX-FLAG-YTHDF1 W465A, and psiCheck2-m^6^A-null-ISRE-*IFITM1* 3’ UTR reporter (wild-type and m^6^A-mut). pLEX-FLAG-METTL14 was generated by cloning the PCR-amplified FLAG-tagged METTL14 coding sequence into the BamHI and XhoI restriction sites of the pLEX expression vector. pLEX-FLAG-YTHDF1 W465A was generated by site-directed mutagenesis of pLEX-FLAG-YTHDF1. WT and m^6^A-mut IFITM1 3’ UTR reporter plasmids (psiCheck2-m^6^A-null-ISRE-*IFITM1* 3’ UTR reporter) were generated by inserting either wild-type *IFITM1* 3’ UTR cDNA or *IFITM1* 3’ UTR cDNA with 4 A-to-G mutations at potential m^6^A sites (obtained as IDT gBlocks) into the XhoI and NotI restriction sites of psiCheck2-m^6^A-null (Gokhale et al., 2019). The 5X ISRE promoter was PCR-amplified from pISREluc (Sumpter et al., 2005) then inserted into the KpnI and NheI sites. All DNA sequences were verified by sequencing.

### Transfection

siRNAs directed against METTL3 (SI04317096), METTL14 (SI00459942), or non-targeting AllStars negative control siRNA (1027280) were purchased from Qiagen. All siRNA transfections were performed using the Lipofectamine RNAiMax reagent (Invitrogen), according to manufacturer’s instructions. siMETTL3/14 co-transfections were performed at a ratio of 1:2 siMETTL3:siMETTL14. Huh7 and A549 cells were transfected with 25 pmol of siRNA at a final concentration of 0.0125 μM, and NHDF cells were transfected with 250 pmol of siRNA at a final concentration of 0.25 μM. Media was changed 4 hours post-transfection, and cells were incubated for 36 h post-transfection prior to each experimental treatment. Plasmid transfections of IFITM1 3’ UTR reporter plasmids (500 ng per single well of a 6-well plate) were performed using the FuGENE 6 (Promega), according to manufacturer’s instructions.

### Generation of Overexpression Cell Lines

Lentiviral particles were generated by harvesting supernatant 72 h post-transfection of 293T cells with pLEX-FLAG-METTL14, pLEX-FLAG-YTHDF1, or pLEX-FLAG-YTHDF1 W465A, and the packaging plasmids psPAX2 and pMD2.G (provided by Duke Functional Genomics Facility). This supernatant was then used to transduce Huh7 cells for 48 hours. Following transduction, cells were selected in 2 μg/mL puromycin (Sigma) for 48 hours and then single cell colonies were isolated. Overexpression of FLAG-tagged proteins in selected colonies was verified by immunoblotting, and we also verified that METTL14 overexpression stabilized METTL3 (Ping et al., 2014), creating METTL3/14 overexpression cell lines. These clones were maintained in cDMEM containing 1 μg/mL puromycin.

### Immunoblotting

Cells were lysed in a modified radioimmunoprecipitation assay (RIPA) buffer (10 mM Tris [pH 7.5], 150 mM NaCl, 0.5% sodium deoxycholate, and 1% Triton X-100) supplemented with protease inhibitor cocktail (Sigma) and phosphatase inhibitor cocktail II (Millipore), and post-nuclear lysates were harvested by centrifugation. Quantified protein (between 5 and 15 μg) was added to a 4X SDS protein sample buffer (40% glycerol, 240 mM Tris-HCl [pH 6.8], 8% SDS, 0.04% bromophenol blue, 5% beta-mercaptoethanol), resolved by SDS/PAGE, and transferred to nitrocellulose membranes in a 25 mM Tris-192 mM glycine-0.01% SDS buffer. Membranes were stained with Revert 700 total protein stain (LI-COR Biosciences), then blocked in 3% bovine serum albumin. Membranes were incubated with primary antibodies for 2 hours at room temperature or overnight at 4°C. After washing with PBS-T buffer (1× PBS, 0.05% Tween 20), membranes were incubated with species-specific horseradish peroxidase-conjugated antibodies (Jackson ImmunoResearch, 1:5000) for 1 hour at room temperature, followed by treatment of the membrane with Clarity enhanced chemiluminescence (Bio-Rad) and imaging on an Odyssey Fc imaging system (LI-COR Biosciences). The following antibodies were used for immunoblotting: mouse anti-IFITM1 (Proteintech 60074-1-Ig, 1:1000; recognizes IFITM1 but not IFITM2 or IFITM3 (Shi et al., 2018; Xie et al., 2015)), rabbit anti-MX1 (Abcam ab207414, 1:1000), mouse anti-ISG15 (Santa Cruz sc-166755, 1:5000), rabbit anti-EIF2AK2 (Abcam ab32506, 1:1000), rabbit anti-METTL14 (Sigma HPA038002, 1:2500), mouse anti-METTL3 (Abnova H00056339-B01P, 1:1000), rabbit anti-YTHDF1 (Proteintech 17479-1-AP, 1:1000), mouse anti-FLAG-HRP (Sigma A8592, 1:5000), rabbit anti-GFP (Thermo Fisher Scientific A-11122, 1:1000).

### Quantification of Immunoblots

Following imaging using the LI-COR Odyssey Fc, immunoblots were quantified using ImageStudio Lite software, and raw values were normalized to total protein (Revert 700 total protein stain) for each condition.

### MeRIP-seq and Analysis

Following mock or IFN-β treatment of Huh7 cells for 8 hours, cellular RNA was harvested using TRIzol (Thermo Fisher Scientific), polyA-tailed mRNA was selected using the Dynabeads mRNA Purification kit (Thermo Fisher Scientific), and MeRIP-seq was performed using the NEB EpiMark m^6^A-enrichment kit as previously described (Gokhale et al., 2019) with the following modifications. Briefly, 25 mL Protein G Dynabeads (Thermo Fisher) per sample were washed three times in MeRIP buffer (150 mM NaCl, 10 mM Tris-HCl [pH 7.5], 0.1% NP-40), and incubated with 1 mL anti-m6 A antibody for 2 h at 4C with rotation. After washing three times with MeRIP buffer, anti-m6 A conjugated beads were incubated with purified mRNA with rotation at 4C overnight in 300 mL MeRIP buffer with 1 mL RNase inhibitor (recombinant RNasin; Promega). 10% of the mRNA sample was saved as the input fraction. Beads were then washed twice with 500 mL MeRIP buffer, twice with low salt wash buffer (50 mM NaCl, 10 mM Tris-HCl [pH 7.5], 0.1% NP-40), twice with high salt wash buffer (500 mM NaCl, 10 mM Tris-HCl [pH 7.5], 0.1% NP-40), and once again with MeRIP buffer. m6 A-modified RNA was eluted twice in 100 mL of MeRIP buffer containing 5 mM m6 A salt (Santa Cruz Biotechnology) for 30 min at 4C with rotation. Eluates were pooled and concentrated by ethanol purification. RNA-seq libraries were prepared from both eluate and 10% input mRNA using the TruSeq mRNA library prep kit (Illumina), subjected to quality control (MultiQC), and sequenced on the HiSeq 4000 instrument. Reads were trimmed using Trimmomatic (Bolger et al., 2014) and aligned to the hg38 genome using the splice-aware STAR aligner (Dobin et al., 2013). Changes in gene expression between Mock and IFN-β treated samples were then identified using DESeq2 (Love et al., 2014) based on differences in read counts from featureCounts (Liao et al., 2014) and plotted in Figure S2A. m^6^A peaks were identified in IFN-β treated samples using the MeTDiff peak caller (Cui et al., 2018) and additionally with meRIPPer (https://sourceforge.net/projects/meripper/). Presented data are from MeTDiff analysis unless otherwise noted. Raw data from Winkler et al. (Winkler et al., 2019) and Rubio et al. (Rubio et al., 2018) were similarly processed (Figure S2B). Coverage plots were generated using CovFuzze (Imam et al., 2018) and a metagene plot for peak locations produced as previously described (Gokhale et al., 2019). Motif enrichment was calculated using HOMER (Heinz et al., 2010). Full methods and scripts for data processing are open-source and online on GitHub (https://github.com/al-mcintyre/merip_reanalysis_scripts) (McIntyre et al., 2020).

### RT-qPCR

Total cellular RNA was extracted using the Qiagen RNeasy kit (Life Technologies) or TRIzol extraction (Thermo Fisher Scientific). RNA was then reverse transcribed using the iScript cDNA synthesis kit (Bio-Rad) as per the manufacturer’s instructions. The resulting cDNA was diluted 1:5 in nuclease-free H2O. RT-qPCR was performed in triplicate using the Power SYBR Green PCR master mix (Thermo Fisher Scientific) and the Applied Biosystems Step One Plus or QuantStudio 6 Flex RT-PCR systems. The oligonucleotide sequences used are listed in Table S5.

### Nuclear/Cytoplasmic Fractionation

Following siRNA treatment (36 h) and IFN-β treatment (20 h), cells were harvested and lysed in 200 μL lysis buffer (10 mM Tris-HCl [pH 7.4], 140 mM NaCl, mM MgCl2, 10 mM EDTA, 0.5% Nonidet P-40 (NP-40)) on ice for 10 minutes. Following centrifugation at 12000 X g at 4°C for 5 minutes, the supernatant (cytoplasmic fraction) was collected, and the nuclear pellet was rinsed twice with lysis buffer. RNA was extracted from cytoplasmic and nuclear pellets using TRIzol reagent and analyzed by RT-qPCR.

### Protein Stability Analysis

Following siRNA treatment (36 h), Huh7 cells were treated with IFN-β for 16 hours to induce ISGs. IFN-β was then replenished at half the dose in cDMEM containing either DMSO as a control, or 50 μg/mL cycloheximide (CHX, Sigma-Aldrich). Cells were harvested over a timecourse (0, 2, 4, 6, 8, 12 hours post-CHX) and subjected to immunoblotting. Protein stability was determined by measuring the protein remaining at each timepoint following CHX treatment.

### Polysome Profiling

Cells treated with siRNAs (36 h) were treated with IFN-β for 6 hours, then pulsed with CHX (50 μg/mL) for 10 minutes. Cells were harvested using trypsin and then lysed in cytoplasmic lysis buffer (200 mM KCl, 25 mM HEPES [pH 7.0], 10 mM MgCl2, 2% n-Dodecyl β-D-maltoside (DDM; Chem-Impex), 0.2 mM CHX, 1 mM DTT, 40 U RNasin) for 15 minutes on ice. Following clarification, lysates were ultracentrifuged on 15-50% sucrose gradients prepared in polysome gradient buffer (200 mM KCl, 25 mM HEPES [pH 7.0], 15 mM MgCl2, 1 mM DTT, 0.2 mM CHX) at 35,000 X g for 3.5 hours at 4°C. Following ultracentrifugation, 24 fractions were collected from each sample using a BioComp Piston Gradient Fractionator instrument fitted with a TRIAX flow cell to measure absorbance. RNA was extracted from each fraction using TRIzol LS reagent (Thermo Fisher Scientific), and RNA quality was checked on a 1% agarose gel. Following cDNA synthesis using the iScript cDNA synthesis kit (Bio-Rad), RT-qPCR was performed using primers specific for each gene.

### MeRIP-RT-qPCR

Total cellular RNA was harvested using TRIzol reagent and normalized to equal input concentrations. m^6^A-positive and m^6^A-negative control oligonucleotides (EpiMark N6-Methyladenosine Enrichment Kit, New England Biolabs) were spiked into total RNA prior to immunoprecipitation. RNA was then immunoprecipitated with anti-m^6^A antibody (New England Biolabs) overnight at 4°C with head-over-tail rotation, and then washed twice with 1X reaction buffer (150mM NaCl, 10mM Tris-HCl, pH 7.5, 0.1% NP40), twice with low salt wash buffer (50 mM NaCl, 10 mM Tris-HCl, pH 7.5, 0.1% NP-40), twice with high salt wash buffer (500 mM NaCl, 10 mM Tris-HCl, pH 7.5, 0.1% NP-40), and once with 1X reaction buffer. RNA was eluted from beads in elution buffer twice for 1 hour at 4°C, and then precipitated in isopropanol overnight at −20°C, pelleted by centrifugation, and resuspended in nuclease-free water. Equal volumes of eluted RNA and input RNA were used for cDNA synthesis and quantified by RT-qPCR. IP efficiency was normalized by relative pulldown of spike-in positive controls.

### Luciferase Assays

Following plasmid transfection of WT and m^6^A-mut IFITM1 3’ UTR reporters and mock or IFN-β treatment (12 h), a dual luciferase assay (Promega) was performed according to manufacturer’s instructions. Data was normalized as fold-change (IFN-β over mock) of the value of *Renilla* luminescence divided by firefly luminescence, and values for WT IFITM1 3’ UTR reporter were set as 1.

### RNA Immunoprecipitation

Following DNA transfection (16 h) and IFN-β treatment (8 h), cells were harvested and lysed in polysome lysis buffer (100 mM KCl, 5 mM MgCl2, 10 mM HEPES [pH 7.0], 0.5% NP-40) supplemented with protease inhibitor cocktail (Sigma) and RNasin ribonuclease inhibitor (Promega), and lysates were cleared by centrifugation. Ribonucleoprotein complexes were immunoprecipitated with anti-FLAG M2 beads (Sigma) overnight at 4°C with head-over-tail rotation, and then washed five times in ice-cold NT2 buffer (50 mM Tris-HCl [pH 7.4], 150 mM NaCl, 1 mM MgCl2, 0.05% NP-40). Protein for immunoblotting was eluted from 25 percent of beads by boiling in 2X Laemmli sample buffer (Bio-Rad). RNA was extracted from 75 percent of beads using TRIzol reagent (Thermo Fisher Scientific). Equal volumes of eluted RNA were used for cDNA synthesis, quantified by RT-qPCR, and normalized to RNA levels in input samples. Enrichment over GFP was then calculated and plotted.

### RNA-seq

Following siRNA treatment (36 h), Huh7 cells seeded in 10-cm^2^ plates were stimulated with IFN-β or mock treated (8 h), then harvested and RNA extraction was performed using TRIzol reagent (Thermo Fisher Scientific). Samples were then treated with Turbo DNase I (Thermo Fisher Scientific) according to manufacturer protocol and incubated at 37°C for 30 min, followed by phenol/chloroform extraction and ethanol precipitation overnight. RNA concentrations were then normalized. Sequencing libraries were prepared using the KAPA Stranded mRNA-Seq Kit (Roche) and sequenced on an Illumina HiSeq 4000 with 100 bp paired-end reads by the Duke University Center for Genomic and Computational Biology.

### Ribo-seq

Following siRNA treatment (36 h), Huh7 cells seeded in 15-cm^2^ plates were stimulated with IFN-β (8 h), then washed with ice cold PBS, and flash frozen in liquid nitrogen. Cells were then lysed in plates with polysome lysis buffer (20 mM Tris-HCl [pH 7.4], 150 mM NaCl, 5 mM MgCl_2_, 1 mM DTT, 1% Triton X-100, 25 U/mL Turbo DNase I (Thermo Fisher Scientific)), scraped, and passed through a 25 gauge needle before collection in microfuge tubes and incubation for 15 minutes on ice. Cytoplasmic lysates were clarified by centrifugation. 5% of lysate was taken for western blotting, and the remaining cytoplasmic lysate was supplemented with 0.4M CaCl_2_ and 4000 gel units micrococcal nuclease (New England Biolabs), and incubated at 37°C (30 min) to generate ribosome protected fragments (RPF RNA). RPF RNA was then ultracentrifuged (35000 X g at 4°C for 3.5 h), over 15-50% sucrose gradients in polysome gradient buffer (20 mM Tris-HCl [pH 7.4], 150 mM NaCl, 5 mM MgCl_2_, 1 mM DTT), after which 12 fractions were collected from each sample using a BioComp Piston Gradient Fractionator instrument fitted with a TRIAX flow cell to measure absorbance. Monosome fractions (fractions 6 and 7) were then pooled and loaded onto a 100 kD molecular weight cut-off filter (Vivaspin 20) and centrifuged at 3000 X g at 4°C for 35 minutes to concentrate monosome-bound RPF RNA. The flow-through was discarded and retained monosomes were separated from RPF RNA by adding polysome lysis buffer supplemented with 50 mM EDTA and incubation on ice for 15 minutes. The resulting RPF RNA solution was then re-applied to the emptied 100 kD molecular weight cut-off filter and centrifuged at 3000 X g at 4°C for 15 minutes to separate RPF RNA from monosomes. The flow-through containing the RPF RNA was then collected, phenol-chloroform extracted, and ethanol precipitated. Precipitated RPF RNA samples were then run on a 15% TBE-Urea gel (Invitrogen), and a band corresponding to 28-32 nucleotides was excised, crushed, and incubated in 0.4M NaCl with 40 units of RNasin ribonuclease inhibitor (Promega) for 8 hours shaking at 4°C 1100 RPM. RNA was recovered by filtration through Corning Costar Spin-X columns (Sigma-Aldrich) then isopropanol precipitated overnight. After resuspension, the remaining RNA was T4 Polynucleotide Kinase (New England Biolabs) treated, phenol-chloroform extracted, and precipitated in ethanol overnight. Sequencing libraries for RPF samples were then generated using the NEB Next small RNA library prep kit and these libraries were sequenced on an Illumina NextSeq 500 High-output 75 bp with paired end reads by the Duke University Center for Genomic and Computational Biology.

### RNA-seq and Ribo-seq Data Analysis

Reads were evaluated using FastQC and trimmed using cutadapt (Martin, 2011), followed by alignment to the hg38 human reference genome using the STAR aligner with default parameters. The number of read fragments uniquely aligned to each gene were counted with the Gencode v21 main comprehensive gene annotation file (aggregated by gene_name) using featureCounts. Using a python script, the raw counts from each replicate and condition were merged to generate a count matrix with N rows/genes and M samples/columns (python scripts for count-matrix generation are open-source and online on GitHub (https://github.com/hmourelatos/McFadden_ISG_m6a_countMatricies)). To identify differentially expressed genes between various groups, we used DESeq2 (Love et al., 2014) to perform three pairwise contrasts. First, with RNA-seq we compared the effects of IFN-β and mock treatment in cells transfected with siCTRL (Table S1.1). Additional RNA-seq analyses included comparison of siMETTL3/14 and siCTRL treated cells after both IFN-β and mock treatment (Tables S1.2 and S1.3, respectively). Finally, with Ribo-seq, we compared siMETTL3/14 and siCTRL treated cells following IFN-β treatment (Table S4). In each case, DESeq2 was applied with no additional covariates and results shown in Tables S1.1, S1.2, and S3 respectively. Metagene plots from Ribo-seq reads were composed using deepTools v3.1 (Ramírez et al., 2016) with the computeMatrix utility. RNA-seq heatmap was generated using R software, and the heatmap for Ribo-seq was generated using ClustVis (Metsalu and Vilo, 2015).

### Mass Spectrometry

Prior to the siRNA experiments, cells were grown for at least 12 generations in DMEM medium without Lysine and Arginine (#PI88420), supplemented with Dialyzed FBS (#26000044), either light or heavy L-Arginine and L-Lysine (L-Arginine-HCl #PI88427; L-Arginine-HCl, 13C6, 15N4 #PI88434; L-Lysine-2HCl #PI88429; L-Lysine-2HCl, 13C6, 15N2 #PI88432), and 100 U/ml penicillin, 100 μg/ml streptomycin and 2 mM L-glutamine. Following stable isotope labeling, siRNA-treated cells (36 h) were stimulated with IFN-β for 24 hours prior to harvest by trypsinization and lysis in RIPA buffer supplemented with protease inhibitor cocktail (Sigma) and phosphatase inhibitor cocktail II (Millipore), and post-nuclear lysates were harvested by centrifugation. 5 μL at 1 μg/μL of siMETTL3/14 (Heavy) extracts were mixed with 5 μL at 1 μg/μL of siCTRL (Light) extracts for the Forward experiment, and 5 μL at 1 μg/μL of siMETTL3/14 (Light) extracts were mixed with 5 μL at 1 μg/μL of siCTRL (Heavy) extracts for the Reverse experiment. The lysates were run on a 4-12% Bis-Tris gel for 30 min. The gel was stained with Colloidal Coomassie and a single patch was cut and processed for each sample. The gel patches were digested with trypsin. The resulting peptides were cleaned with a C18 tip. Liquid chromatography was performed with an EASY-nLC™ 1000 Integrated Ultra High Pressure Nano-HPLC System and MS/MS with a Q-EXACTIVE System equipped with a Nanospray Flex Ion Source, as previously described (Abell et al., 2017).

### Mass Spectrometry data analysis

Four RAW files representing two replicates each of Forward and Reverse SILAC experiments were retrieved from the Orbitrap. Heavy/light label ratios were quantified across all samples using MaxQuant v1.6.7.0 with the Andromeda search engine and default parameters other than specifying SILAC labels (Cox and Mann, 2008; Cox et al., 2011). For all analyses, the “H/L Ratio – Normalized” field containing median-centered label ratios was extracted for each peptide and/or protein and compared across replicates (Table S3). Heatmaps for mass spectrometry were generated using ClustVis (Metsalu and Vilo, 2015).

#### Peptide regression modeling

To take advantage of the measurement independence of unique peptides, we applied a simple linear mixed model to identify significant shifts in labeling ratio between conditions while accounting for peptide-specific effects. First, we merge all proteins that are described by the same set of peptide ratios (e.g. protein sequences from the same gene for which all detected peptides are shared). Then, for each protein (defined now as a set of peptide ratios), we fit a linear model of the following form using lme4 in R (Bates et al., 2015)

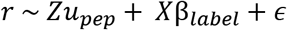

where:

- *r* is the median-normalized heavy/light label ratio derived from MaxQuant
- *Z* is a binary design matrix indicating the peptide identity of each ratio measurement
- *X* is a binary design vector indicating condition (forward or reverse)
- *u_pep_* is a vector of random effects corresponding to each peptide effect
- β_*label*_ is the fixed effect of condition

Thus, each peptide ratio is described as the sum of a peptide-level random effect and a condition (forward vs reverse) fixed effect, and some error. We extract effect size estimates and p-values from unmodified Wald tests on the fixed effect of condition, and adjust across all proteins with the Benjamini-Hochberg (BH) procedure. Note that in the less-powerful case of proteins with only one measured peptide, the random peptide effect is just a constant and the model reduces to simply comparing the means of the forward and reverse replicate ratios for the single peptide.

#### Aggregating proteins for gene-level results

Peptide regression modeling generates one test for each protein, so many genes are tested multiple times at each of their proteins. Annotated reference protein sequences often contain multiple entries per gene with varying degrees of similarity. After applying the procedure above, we observe as expected that the vast majority of genes contain either all significant or all non-significant protein results. We conservatively describe as significant any gene with a significant maximum p-value (meaning all tested proteins are significant) following multiple test correction.

### Lead Contact and Materials Availability

Further information and requests for resources and reagents should be directed to and will be fulfilled by the Lead Contact, Stacy M. Horner.

### Data Availability

All raw data from RNA-seq, MeRIP-seq, and Ribo-seq are available through GEO (accession number: GSE155448).

Raw data from mass spectrometry are available at the following URLs:

https://web.corral.tacc.utexas.edu/xhemalce/Forward1.raw
https://web.corral.tacc.utexas.edu/xhemalce/Forward2.raw
https://web.corral.tacc.utexas.edu/xhemalce/Reverse1.raw
https://web.corral.tacc.utexas.edu/xhemalce/Reverse2.raw

## Supplemental Information

Supplemental information Figures S1-S4.

Table S1: RNA-seq analysis of gene expression changes following IFN-β treatment and METTL3/14 depletion.

- Table S1.1: siCTRL IFN / siCTRL Mock
- Table S1.2: siMETTL3/14 Mock / siCTRL Mock
- Table S1.3: siMETTL3/14 IFN / siCTRL IFN

Table S2: m^6^A peaks in the IFN-β induced transcriptome.

- Table S2.1: Input RNA-seq Analysis (IFN / Mock)
- Table S2.2: MeTDiff m^6^A Peaks
- Table S2.3: meRIPPer m^6^A Peaks

Table S3: Analysis of METTL3/14 depletion effect on protein expression using quantitative mass spectrometry.

Table S4: Analysis of Ribo-seq data from METTL3/14 depletion and IFN-β treatment.

Table S5: List of oligonucleotides and siRNAs used in this study.

