## Supplemental Figures for "Post-Transcriptional Regulation of Antiviral Gene Expression by *N6*-Methyladenosine"

**SUPPLEMENTAL INFORMATION**

**Table of contents:**

Figure S1: Related to Figure 1.

Figure S2: Related to Figure 1.

Figure S3: Related to Figure 2.

Figure S4: Related to Figure 5.

**Figure S1: Related to Figure 1.**

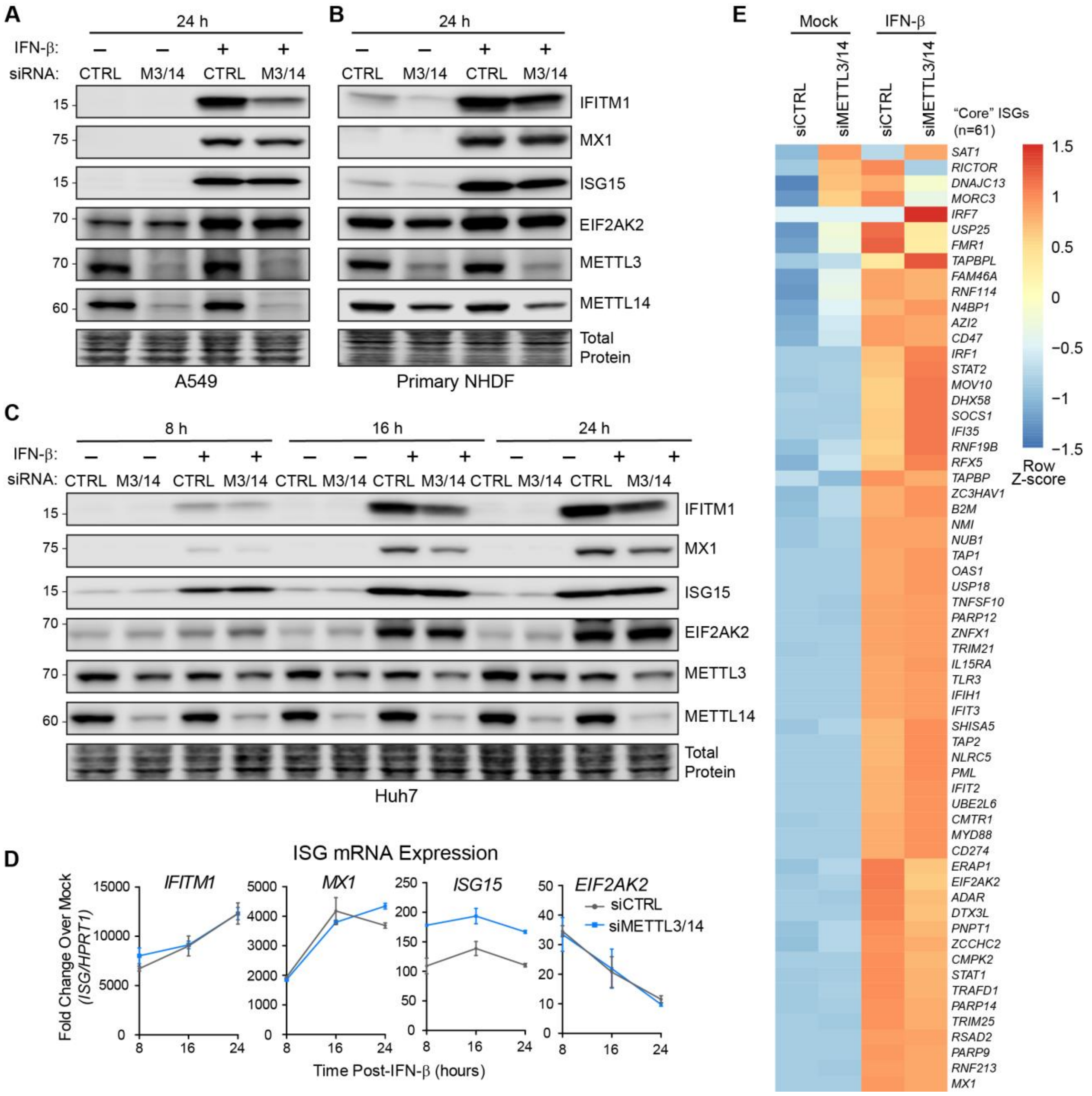


**Figure S1: Related to Figure 1. (A-C)** Immunoblot analysis of extracts from A549 cells (A), primary neonatal human dermal fibroblast cells (B), or Huh7 cells (C) transfected with siRNAs to METTL3/14 (M3/14) or control (CTRL) (36 h) prior to treatment with mock or IFN-β for 24 h (A, B) or a timecourse of 8 hours, 16 hours, and 24 hours (C). All immunoblots are representative of 4 biological experiments. **(D)** RT-qPCR analysis of ISG induction normalized to *HPRT1* following IFN-β treatment of Huh7 cells treated with non-targeting control (CTRL) or METTL3/14 siRNA plotted as fold change over mock treatment for each ISG. Values are the mean ± SD of 3 technical replicates, representative of 4 experiments. **(E)** RNA-seq analysis following siRNA transfection and IFN-β treatment (3 biological replicates). Heat map shows the Row Z-score for each ISG (n=61) with Euclidean row clustering.

**Figure S2: Related to Figure 1.**

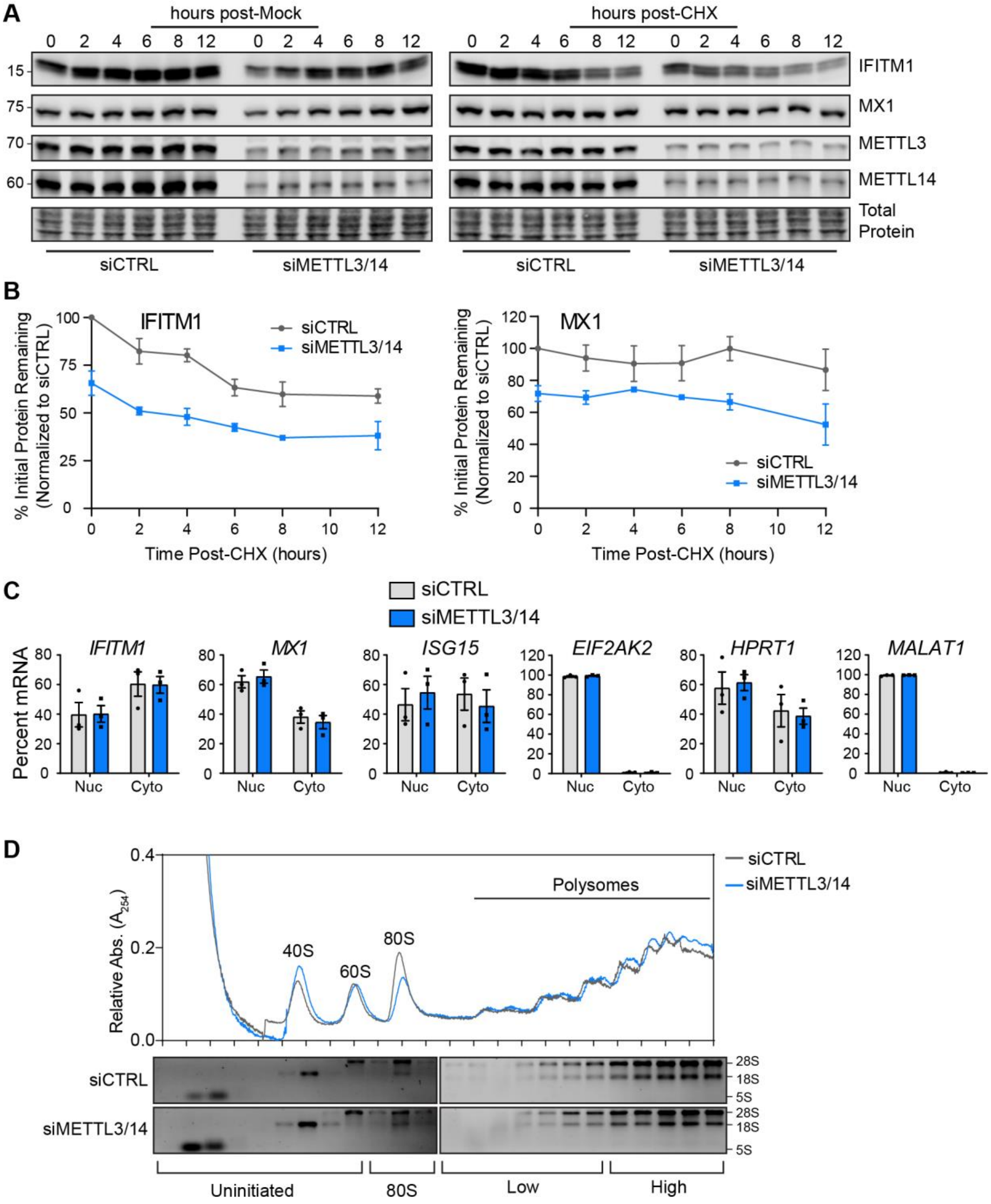

**Figure S2: Related to Figure 1. (A)** Immunoblot analysis of ISG protein levels following cycloheximide (CHX) treatment. Following siRNA transfection, cells were treated with IFN-β (16 h) to induce ISGs, then mock or CHX treated along with a second dose of IFN-β, and lysates were harvested at indicated timepoints. **(B)** Quantification of ISG expression in (A), normalized to total protein, and plotted as percent of initial protein remaining at each timepoint, relative to siCTRL timepoint 0. **(C)** RT-qPCR analysis of the percent mRNA in nuclear (Nuc) and cytoplasmic (Cyto) fractions in siRNA-treated Huh7 cells following IFN-β (12 h). **(D)** Relative absorbance values of sucrose gradient fractions isolated from extracts of IFN-β-treated (6 h) Huh7 cells treated with indicated siRNA. The 40S, 60S, and 80S subunit peaks, as well as polysome peaks, are noted. RNA from each fraction was separated by agarose gel electrophoresis and visualized with ethidium bromide to see the ribosomal RNA bands. Values are the mean ± SEM of 3-4 biological replicates (B), and the mean ± SEM of 3 biological replicates (C). Data in (D) are representative of 3 biological experiments.

**Figure S3: Related to Figure 2.**

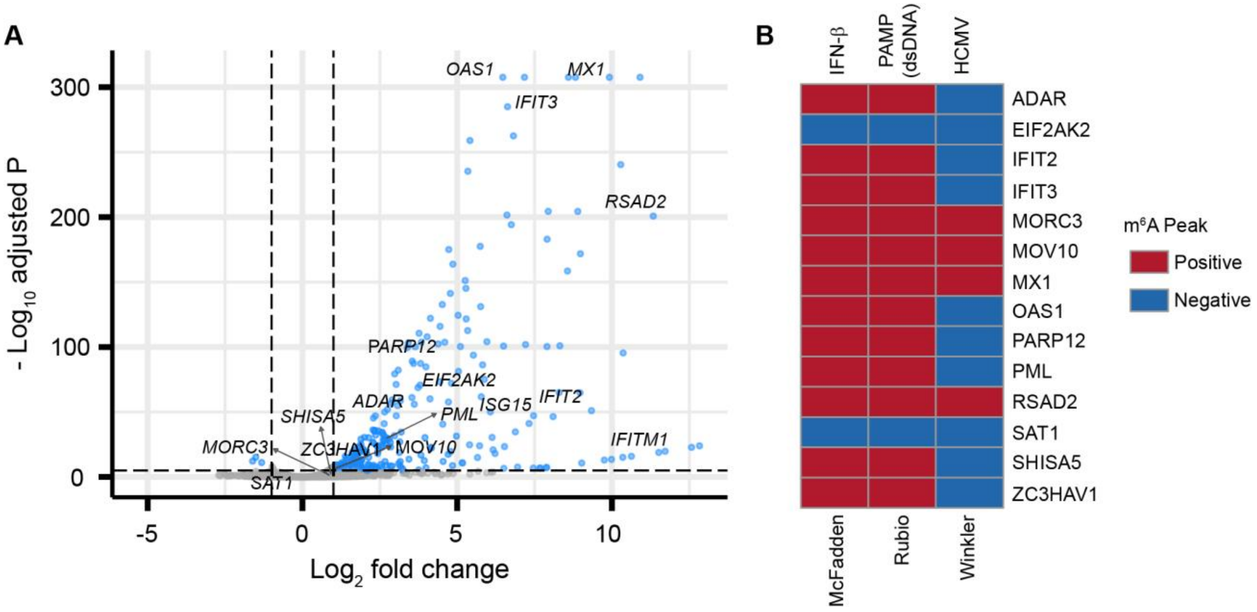


**Figure S3: Related to Figure 2. (A)** Volcano plot showing gene expression changes (58,037 total genes) following IFN-β treatment (8 h) compared to mock treatment, with core antiviral ISGs labeled. Analysis was performed on 3 biological replicates of each condition. **(B)** Analysis of MeRIP-seq data from this study and two previous studies (Rubio et al., 2018; Winkler et al., 2019) showing $m^6A$ peak presence or absence in core antiviral ISGs following induction of IFN pathways. IFN responses were induced by IFN-β (McFadden), double stranded DNA (Rubio), or HCMV (Winkler).

**Figure S4: Related to Figure 5.**

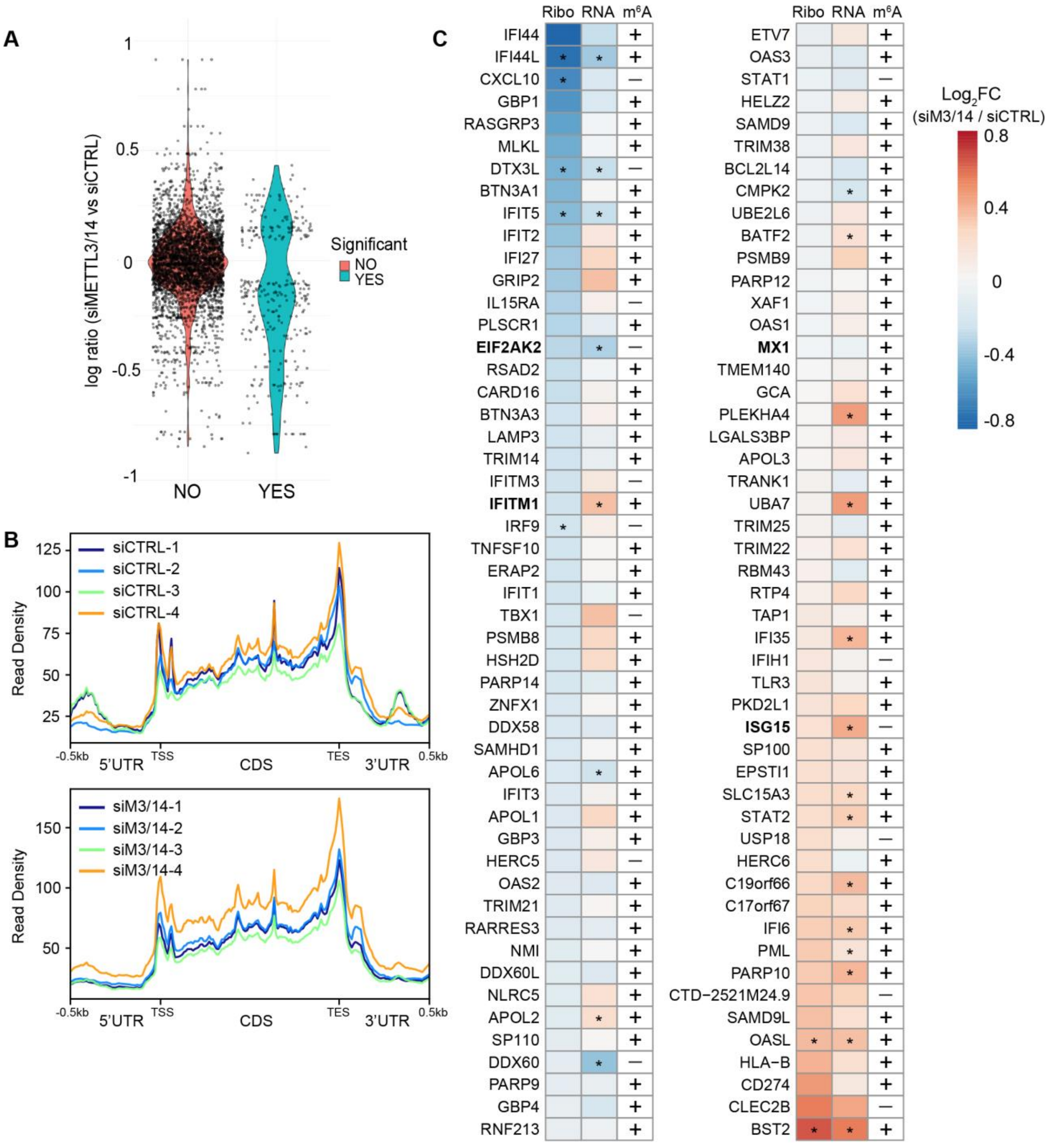


**Figure S4: Related to Figure 5. (A)** Violin plot of the effect of siMETTL3/14 treated Huh7 cells compared to siCTRL treatment (+ IFN-β 24 h; 2 biological replicates each), expressed as log ratio, separated in two groups: non-significant (NO) with an adjusted p value > 0.01, or significant (YES) with an adjusted p value < 0.01. **(B)** Metagene plot showing ribosome protected fragment abundance 500 base pairs upstream and downstream of the translation start site (TSS) and translation end site (TES), and along the open reading frame (CDS; scaled to 1000 bp) for all siCTRL IFN and all siMETTL3/14 IFN replicates (+ IFN-β 8 h; n=4 biological replicates). **(C)** 3-column heatmap shows the effect of METTL3/14 depletion on the expression of 100 ISGs. The first column shows the $\log_2$fold change of Ribo-seq reads (siMETTL3/14 over siCTRL). The second column shows $\log_2$ fold change of mRNA reads from an independent RNA-seq experiment (siMETTL3/14 over siCTRL), and the third column indicates $m^6A$ status (+ indicates $m^6A$-positive; - indicates $m^6A$-negative) from MeRIP-seq. Genes include the top 100 most highly-induced genes by IFN treatment from RNA-seq (see also Table S1) with base mean > 25, as determined by DEseq2 (Love et al., 2014) in all conditions (Ribo-seq, RNA-seq, MeRIP-seq). ISGs investigated in other figures are shown in bold. * adjusted P < 0.05.
